# Explainable Deep-Learning on condition specific expression profiles reveals critical cytosines in gene regulation

**DOI:** 10.1101/2025.06.01.657178

**Authors:** Veerbhan Kesarwani, Anchit Kumar, Akanksha Sharma, Sagar Gupta, Kirti Pandey, Gaurav Zinta, Ravi Shankar

**Affiliations:** Studio of Computational Biology & Bioinformatics, The Himalayan Centre for High-throughput Computational Biology, (HiCHiCoB, A BIC supported by DBT, India), Biotechnology Division, CSIR-Institute of Himalayan Bioresource Technology (CSIR-IHBT), Palampur (HP), 176061, India; Integrative Plant AdaptOmics Lab (iPAL), Biotechnology Division, CSIR-Institute of Himalayan Bioresource Technology (IHBT), Palampur, Himachal Pradesh, India; Academy of Scientific and Innovative Research (AcSIR), India

**Author notes:** Authors’ email addresses: VK, AK, AS, SG, KP, GZ.

**Keywords:** Epigenomics, DNA methylation, deep learning, ResNet, critical cytosines, gene regulation

## Abstract

Compared to other nucleotides, the cytosines stand as the most expressive one for gene regulation in plants due to its status as methylation-based epigenetic switch. Methylation of some of these cytosines may have higher impact on downstream genes, making them critical ones. To this date not much has been done to decipher the criticality of such cytosines. This is first such pioneering study in decoding the critical ‘C’s where a large volume of bisulfite and RNA-seq data, including 232 WGBS and 260 corresponding RNA-seq datasets from *A. thaliana* and rice, have been utilized. Using a deep learning system, strong relationship was established between methylation states of cytosines in contextual manner with respect to the downstream genes expression levels. Using the same system, all the 2kb upstream regions for *Arabidopsis* and rice genes have been annotated for critical cytosines. Experimental validation demonstrated specific methylation changes at critical cytosines under heat stress, affecting gene expression. The universal method developed may be applied to annotate other plant genomes for critical cytosine identification. GC% similarity, rather than homology, explains gene regulatory behavior under methylation control. The resource, CritiCal-C portal, is publicly accessible, paving the way for revolutionary strategies in gene regulation and expression with minimal interventions.

## Introduction

Epigenetic mechanisms play a vital role in modulating diverse physiological and developmental processes by regulating gene expression and modifying the chromatin accessibility **[1–3]**. Epigenetic marks include chemical modifications of DNA and various post-translational alterations in histone proteins. In plants, DNA methylation has emerged as the most prominent epigenetic regulatory mode, where methylation of cytosines stands out **(Fig. 1a).** This occurs in both symmetric sequences (CG and CHG) and asymmetric sequences, such as CHH (where H represents A, T, or C).

**Figure 1.**
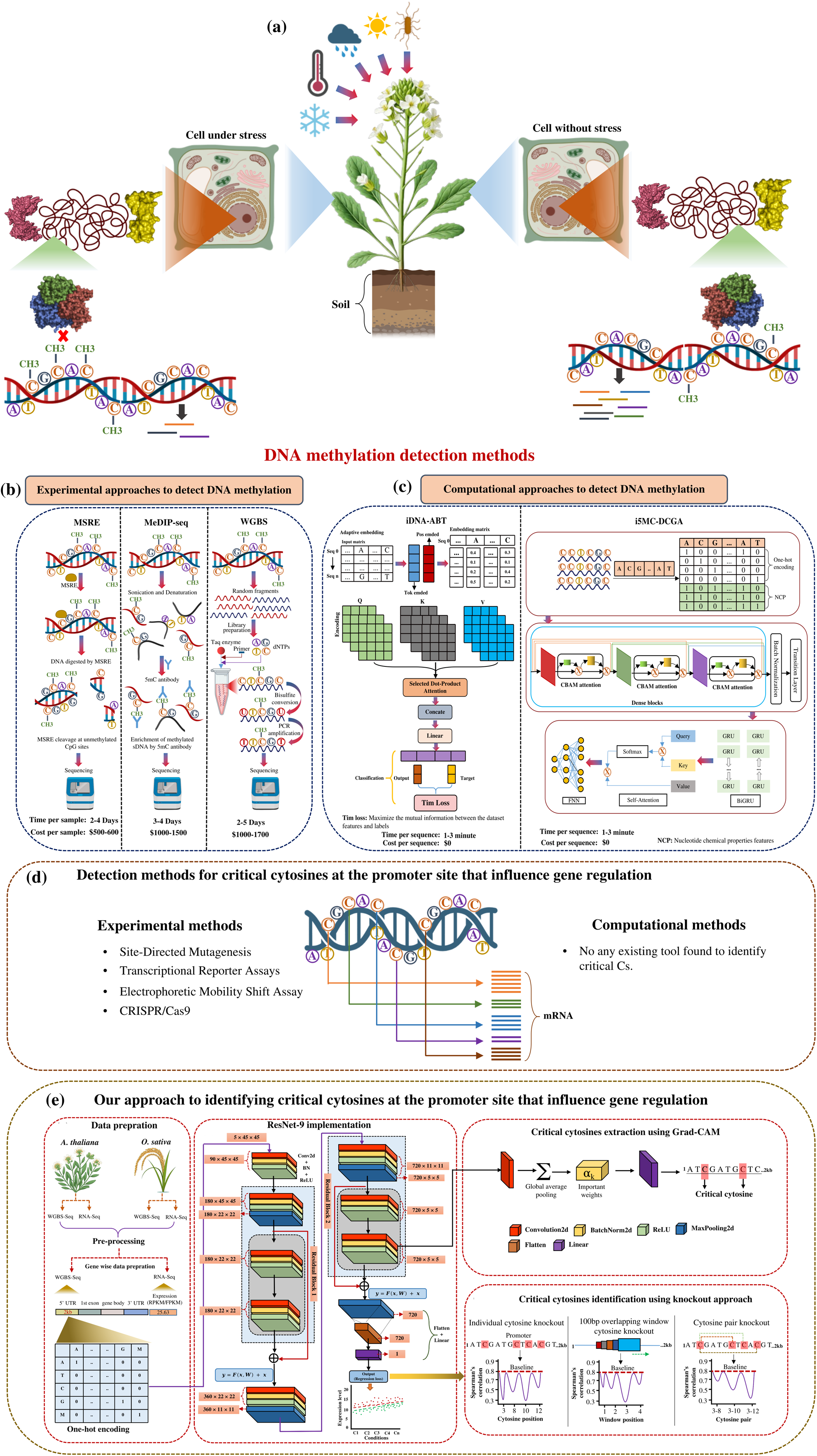
Study Overview. **(a)** Stress-dependent regulation of gene expression by DNA methylation; Right: Under normal conditions, unmethylated DNA permits transcription factor (TF) binding, enabling gene expression, Left: Under biotic/abiotic stress, DNA methylation (red circles) inhibits TF binding, suppressing gene expression. **(b)** Experimental methods for DNA methylation detection. **(c)** Computational approaches for methylation detection. **(d)** Limitations in studying single-cytosine effects, current experimental methods can assess individual cytosine impact but are costly, time-consuming, and condition-specific. No computational tools yet exists to predict expression-level effects of specific cytosines. **(e)** Our solution: Critical cytosine analysis, pipeline for identifying and validating individual cytosines which directly influences gene regulation, bridging the gap between methylation status and expression outcomes.

Despite its significance, the intricate relationship between cytosine and its influence on gene expression across various biological conditions remains poorly understood **[4]**. Promoters host certain regulatory regions located at various intervals from the transcription start site (TSS) controlling the downstream gene expression **(Fig. 1a)** in response to prevailing conditions **[5]**. Cytosine-rich sequences like GC-rich promoters, are frequently associated with strongly controlled transcriptional activity as they regulate the formation of open chromatin structures. Cytosine-rich motifs, such as the G-Box (CACGTG) and ABA-responsive elements (ABRE), are critical for binding transcription factors (TFs) that mediate responses to light, hormones, and stress **[6,7]**. For the reported motifs for various TFs binding sites in *Arabidopsis* at JASPAR database, in ∼60% of the cytosine is found as the most conserved nucleotide. This all places high regulatory stake on cytosines, further amplified by their methylation status.

Various techniques are available to identify methylated regions **(Fig. 1b),** primarily stationed around DNA bisulfite treatment. It converts unmethylated cytosines to uracil, which is then converted to thymine during PCR amplification, and enables the differentiation between methylated and unmethylated cytosines **[8]**. Following bisulfite conversion, one can assess the overall DNA methylation status using microarray-based hybridization techniques such as Illumina HumanMethylation450 (HM450K) and advanced next-generation sequencing (NGS) methods such as whole-genome bisulfite sequencing (WGBS) **(Fig. 1b) [9,10]**. Despite these advancements, the study of DNA methylation remains challenging due to high costs, labor-intensive nature, and technical limitations of these methods, rendering them a costly affair to employ at widescale and frequent uses. **Supplementary File 1** contains detailed information on the advantages and disadvantages of each such technique. Therefore, developing an efficient computational approach to accurately profile DNA methylation under various conditions has become essential.

To tackle these challenges, researchers are increasingly leveraging computational methods, particularly machine learning (ML) and deep learning (DL)-based approaches. Although these tools **(Fig. 1c)** have significantly advanced the field, they are limited in several key aspects. First, they rely on data that are not condition-specific and fail to account for the dynamic nature of DNA methylation across different biological contexts. Secondly, they don’t provide insights into whether the identified methylated cytosines influence gene expression or the specific cytosines that are critical for transcriptional regulation. Very recently, our group developed the DNA Methylation Recognition Unit (DMRU) **[11]** which demonstrated that condition-specific DNA methylation patterns can be accurately identified by integrating transcriptome profiles with 2kb upstream DNA sequence information using deep learning. DMRU successfully leverages gene expression dynamics to propose methylation states across different conditions, achieving >90% accuracy in a cross-species universal manner. However, while DMRU addresses the challenge of identifying where methylation occurs under specific conditions (forward direction: expression → methylation), it does not address the reverse direction: how methylation patterns influence downstream gene expression. Understanding this reverse relationship, from methylation patterns to expression outcomes, remains an essential next step for developing targeted epigenetic interventions. Currently, no computational tool exists to identify which specific cytosines critically influence downstream gene expression. At present, this can only be achieved through experiments such as knocking-out site-directed mutations like cytosine base editors (CBEs) and gene targeting (GT) via homologous recombination (HR) **[12–15].** Performing such experiment for all genes within a species itself is highly impragmatic, expecting them to use widely across species and conditions remains a challenge. Same time, giving criticality weight to cytosines would open several gateways for regulatory interventions, making it highly important to devise some computational approach to obtain it.

To address this problem, as the pioneering work, a novel deep-learning based approach, CritiCal-C, is being presented here (**Fig. 1e**). It utilized 232 WGBS-seq and 260 RNA-seq datasets from *A. thaliana* and *O. sativa*. CritiCal-C has been designed to elucidate the relationship between methylation states in the promoter regions (2kb upstream) and corresponding expression levels of the downstream gene under various environmental conditions. It can identify critical cytosines using three approaches: (1) single cytosine knockout, (2) knockout of each cytosine from a 100 bp overlapping window, and (3) knockout of cytosine base pairs. This approach not only helps to suggest the impactful cytosine methylation points in terms of impact on gene expression but also provides an adaptable and cost-effective alternative to labor-intensive experimental methods such as site-directed mutagenesis, transcription reporter assays, and CRISPR/Cas9-based editing **[16–19] (Fig. 1d)**.

It also functions in a cross-species manner, enabling good generalization of the findings across different plant species, without necessitating species-specific models. Furthermore, a user-friendly web server has been established which would enable the researchers to input methylated promoter sequences of gene of interest to obtain the downstream gene’s expression level, and same time suggesting the critical cytosines within the input promoter sequence. This integrated platform represents a significant advancement in the application of deep learning to plant regulomics, offering an interpretable framework to identify regulatory cytosines.

## Materials and methods

### Data retrieval and pre-processing

This study utilized whole-genome bisulfite sequencing (WGBS-seq) and RNA-seq data obtained from the NCBI Sequence Read Archive (SRA). The dataset comprised 120 WGBS and 135 RNA - seq samples for *A. thaliana*, encompassing 45 unique conditions, and 112 WGBS and 125 RNA-seq samples for *O. sativa*, covering 40 distinct conditions **(Supplementary File 2, Sheet 1-4)**. The selection process for WGBS-seq FASTQ files was contingent on the availability of RNA-seq FASTQ files under the same experimental conditions. Only WGBS-seq files meeting this criterion were used. Genome annotation information was retrieved from Ensembl Plants for both species.

For refinement of both WGBS and RNA-seq data, Trimmomatic v0.39 **[20]** and filteR **[21]** were employed. The processed WGBS reads were subsequently aligned to the reference genomes of both species using Bismark v0.22.2 **[22]**, retaining only uniquely aligned reads and discarding ambiguous mappings. Samtools v0.1.9 **[23]** was used to arrange the reads by genomic coordinates, eliminate PCR duplicates, and convert the processed SAM files to BAM format. The resulting BAM files underwent methylation extraction using Bismark’s methylation extractor function, excluding reads with conversion rates below 90% or fewer than three non-converted cytosines in non-CpG contexts. 2kb upstream promoter sequences for each gene in both species were extracted from the downloaded genome. The downstream gene expression profiles corresponding to each promoter sequence were incorporated into the dataset.

For RNA-seq data, filtered reads were aligned to the reference genomes of both species using HISAT2 **[24]**. Read counting was performed using Rsubread **[25]**. The read counts were subsequently normalized to fragments per kilobase per million mapped reads (FPKM) for paired-end reads and reads per kilobase per million mapped reads (RPKM) for single-end reads, for individual genes across various conditions. For training and testing, the dataset comprising promoter sequences and their corresponding gene expression profiles were partitioned into 70% training set and 30% testing set. In this entire study, for every dataset, it was ensured that no overlap or redundancy existed across training and testing datasets.

### Feature encoding and model construction

One-hot encoding was employed to convert DNA sequences into numerical tensors. The encoding method allocated a unique binary vector to each base state, with A, C, G, T, and M represented as (1,0,0,0,0), (0,1,0,0,0), (0,0,1,0,0), (0,0,0,1,0), and (0,0,0,0,1), respectively. The ‘M’ symbol denotes a methylated cytosine state. The input vector length was 2025 bases, with each base subsequently transformed into a five-dimensional vector, resulting in the vectorization of each sequence into a matrix with dimensions of 2025 × 5 (input sequence length × number of alphabets). Each input sequence was reshaped into a 5-channel tensor, subsequently structured into dimensions of 5 × 45 × 45, representing the spatial dimensions of the input array. The encoded data were modeled through various machine learning and deep-learning approaches. Details about them are provided in **Supplementary File 2, Sheet 5**.

In the present study, ResNet-9 emerged as the most suitable approach among all compared methods for efficiently identifying gene expression. The ResNet-9 architecture **(Fig. 2)** was structured into four distinct stages: convolutional blocks, residual blocks, max-pooling and flattening, and a fully connected layer.

**Figure 2.**
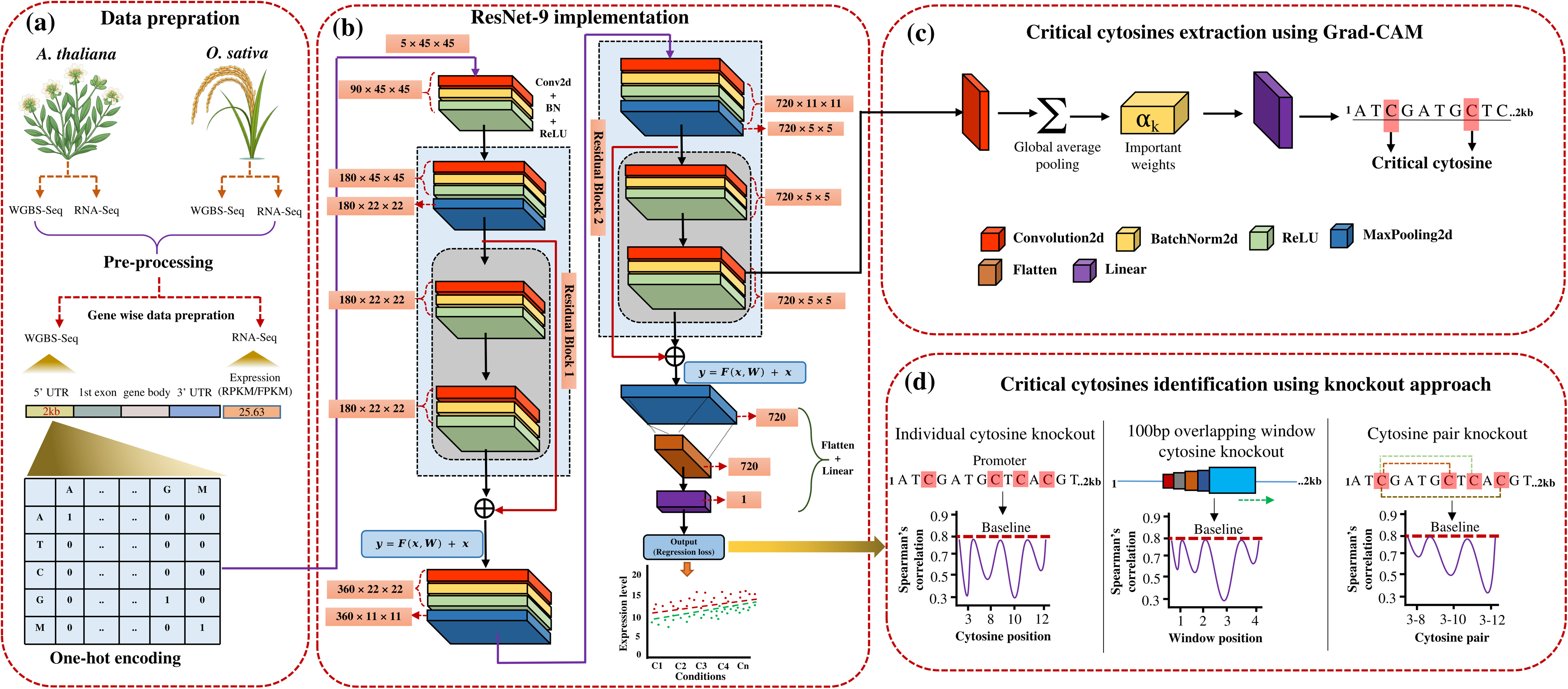
A deep learning-based framework for expression prediction and critical cytosine identification using Grad-CAM and knockout approach. **(a)** The data preparation workflow includes WGBS-Seq and RNA-Seq datasets from *A. thaliana* and *O. sativa* under multiple conditions. For each experimental condition, gene-specific promoter sequences (2kb upstream) are processed by extracting methylation patterns and corresponding gene expression (FPKM/RPKM) levels, which are subsequently transformed into one-hot encoded representations. **(b)** Implementation of the ResNet-9 deep learning model. The architecture processes 5×45×45 input tensors representing the one-hot encoded promoter sequences. Residual blocks enhance feature learning, and the final output is a regression score representing predicted expression. **(c)** Critical cytosines are extracted using Grad-CAM. Feature importance weights are computed and mapped back to the input promoter sequence to identify cytosines that significantly contribute to gene expression prediction. This integrated approach enables systematic identification of functional cis-regulatory elements across species and conditions. **(d)** Critical cytosines are identify using a knockout analysis. Three strategies are illustrated: individual cytosine knockout, 100 bp overlapping window knockout, and cytosine pair knockout. The effect of each knockout on model performance is quantified by changes in Spearman’s correlation with baseline predictions.

#### Convolutional blocks

The network comprised four convolutional blocks (kernel size=3, stride=1). Each block executed a 2D convolution operation, followed by batch normalization and ReLU activation. Max pooling was applied subsequent to selected convolutional layers to reduce the spatial dimensions of the feature maps. The sequence of operations within a convolutional block was as follows:

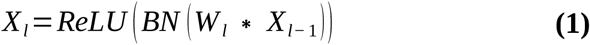

Here, “*X_l_*” is the output feature map of the current layer, *“ X_l−_*_1_ *”* input feature map to the current layer, and *“W _l_***”** convolutional filter/kernel weights for layer.

#### Residual Blocks

Two residual blocks were incorporated into this architecture to mitigate the vanishing gradient problem and enhance the learning efficiency of the model. A residual block establishes a skip connection that enables the network to bypass certain convolutional operations, facilitating the direct transmission of information from earlier layers to later layers. The operation of a residual block can be mathematically defined as:

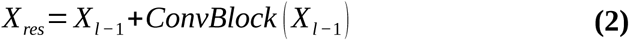

Here, *“ X_l−_*_1_ *”* is the input to the residual block and *“ConvBlock* (*X_l−_*_1_) *”* represents the output from the corresponding convolutional block for that layer.

#### Max Pooling and Flattening

Subsequent to the final convolution block, which encompasses the residual blocks, a max pooling operation was implemented to further reduce the spatial dimensions.

Max pooling selectively retains the most salient features from each spatial region. The resultant output tensor is subsequently flattened into a one-dimensional vector:

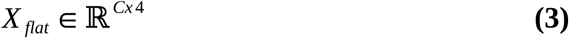

Here, *“ Χ_flat_ ”* is flattened matrix, *“* ℝ*^Cx^* ^4^ *”* represents the matrix in the space of real numbers with shape, *“C ”* is the feature set.

#### Fully Connected Layer

The flattened vector is subsequently processed through a fully connected (dense) layer. This layer maps the high-level features extracted by the convolutional and residual blocks onto a scalar regression output. The output is defined as:

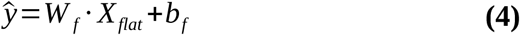

Here, 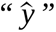 is the estimated regression value, *“W _f_ ”* represents the learned weights for the fully connected layer, and *“ b_f_ ”* is the bias.

In this study, PyTorch was used to implement the CNN, ResNet and DenseNet architectures.

#### Performance measures

To assess the reliability of the predictive models, statistical performance measures like the Spearman’s correlation, RMSE, and MAE were employed. These metrics provide distinct perspectives on model performance: Spearman’s correlation assesses the strength and direction of the relationship between predicted and actual values, whereas RMSE and MAE measure the magnitude of prediction errors. To enhance the reliability, 10-fold random trials of training-testing was employed. The selection of these specific measures facilitated a comprehensive evaluation of the models across various prediction aspects.

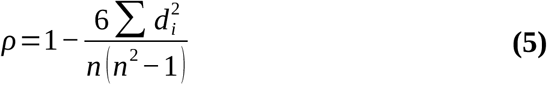

Here, *“ ρ”* Spearman’s rank correlation coefficient (ranges from -1 to +1), *“ d_i_”* difference between the ranks of the two variables, *“ n”* number of observations.

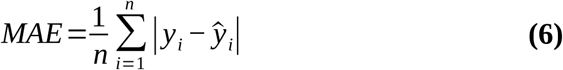

Here, *“ n”* is number of data points, *“ y_i_”* is predicted value, 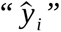 is observed value, 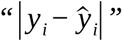 is absolute error.

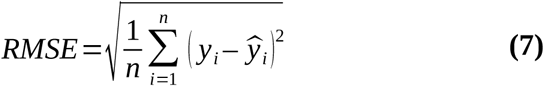

Here, *“ n”* is number of data points, *“ y_i_”* is predicted value, 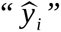 is observed value, 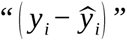 is squared error.

### Hyperparameter Optimization

To optimize the CNN, ResNet and DenseNet-121 architectures, a Bayesian optimization approach was employed using Optuna **[26]**. The optimization process encompassed key architectural and training variables, including dropout (0.1–0.5), learning rate (1e-4 to 1e-1), batch size (1–47), weight decay (1e-6 to 0.1), momentum (1e-5 to 0.5), and epochs (50–100).

### Critical cytosine identification and feature importance determination

To determine the critical cytosines which may influence gene expression, two complementary approaches were employed to examine the input promoter sequences and associated genes from both species. Each method offers distinct insight into the role of cytosine in gene regulation. The first one had three ways: (i) individual cytosines were systematically knocked out from the gene-wise dataset. Following each knock out, the effect of this alteration on the model’s performance in identifying gene expression was evaluated. (ii) 100 base pair (bp) overlapping windows to scan the sequence. All cytosines were knocked out together within each window, analyzing the effect of their absence in every 100 bp segment. The use of 100 bp overlapping windows in DNA sequence analysis enabled to capture the cytosines collectively. (iii) Here, the focus was on identifying the cytosine pairs. The influence of these pairs on gene expression was examined, with particular attention given to pairs whose knockout led to a significant decrease in the Spearman’s correlation.

In the second approach, two different deep-learning based ways were employed to extract critical features. Initially, a custom script were developed to extract weights associated with each cytosine from the eighth convolutional layer. By analyzing these weights, it was possible to quantify the importance of each cytosine in the decision-making process of the model. The second way applied explainability through Grad-CAM **[27]** involving the eighth convolutional layer of the ResNet-9 architecture. The process begins by computing the gradient 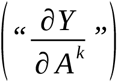 of the resulted gene expression “Y” with respect to the activation maps *“A^k^”* of the eighth convolutional layer. Global average pooling was applied to these gradients to obtain the importance weights 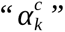 for each channel “c” in the activation map:

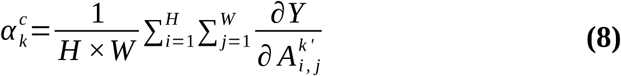

Where, “*H”* and “*W”* are the spatial dimensions of the activation map, and 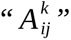 represents the activation at spatial location (*i, j*) in the k-th layer. These weights reflects the contribution of each channel to the prediction.

The Grad-CAM score *“ A^k^ ”* is generated by combining the activation maps ***“A^k^”*** using the importance weights 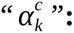

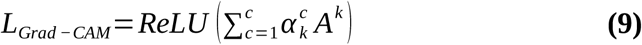

The ReLU function ensured that only positive contributions to the prediction are considered, highlighting the most critical cytosines’ of the input sequence.

### Experimental validation of critical cytosines

#### Plant materials and growth conditions

The wild-type Columbia (Col-0) ecotype *Arabidopsis thaliana* was used. Plants were grown under long-day conditions at 22°C in growth chambers. After 21 days, they were subjected to heat stress at 38°C. Sampling was done after 3 days of heat stress.

#### RNA extraction and quantitative RT-PCR

The RNeasy Plant Mini Kit (74904; QIAGEN) was used to isolate total RNA in accordance with the manufacturer’s instructions. The cDNA was synthesized by 1 μg of DNAse-treated total RNA (Verso cDNA Synthesis Kit: AB-1453/A; thermoscientific). The PCR was setup by mixing 5 μl SYBR Green (Invitrogen,USA), 1 μl cDNA, 0.5 μl forward primer (10 pm/μl), 0.5 μl reverse primer (10 pm/μl), and milli-Q water up to 10 μl and run with cycling conditions 95°C for 3 min followed by 95°C for 30 sec, 60°C for 30 sec, and 72°C for 20 sec, with a melting curved detected at 95°C for 1 min and 65°C for 1 min using CFX96 (Bio-Rad Laboratories). The threshold cycle (Ct) value for each gene was quantified and normalized by Ct value of internal control gene, Actin. The relative expression was calculated using 2^-ΔΔCt^ method converted to log2 fold (Livak and Schmittgen, 2001). Student’s t-test was used as test of significance.

#### Targeted bisulfite sequencing

About 1 μg of genomic DNA from ambient and 38°C treated plants from both the *Arabidopsis thaliana* (Col0) was bisulfite converted using EZ DNA Methylation-Gold™Kit (ZYMO, USA). The bisulfite converted DNA was desalted and 2 μl of it was used for PCR amplification of the selected genomic region representing DMR using locus-specific forward and reverse primers. The purified PCR products were cloned in pGEM-T Easy vector (Promega, USA) and confirmed clones were sequenced via Sanger sequencing method. An average of six clones was chosen randomly for sequencing. Methylation status of cytosines were assessed by comparing the sequence of the ambient and high temperature samples **[28].**

#### Selection of validation targets

AT1G20400 (DUF1204) was selected based on: (1) presence of critical cytosines with post-knockout Spearman correlations <0.45 within a 100 bp promoter window, (2) availability of an adjacent non-critical control region with post-knockout correlations >0.70, and (3) identified stress responsiveness. Two 100 bp windows underwent analysis: the critical window (794-894 bp) contains cytosines showing significant effects on gene regulation in both single and pairwise knockout analyses, while the non-critical window (0-100 bp) contains cytosines exhibiting negligible effects in all knockout analyses. The experimental condition (38°C for 3 days) was not represented in the training dataset, ensuring independent validation.

## Results and Discussion

### Data collection and primary vision of relationship between methylation and gene expression

This study aimed to explore the interplay between the impact of cytosine methylation and gene expression patterns, with a specific emphasis on assessing the important cytosines at different locations which significantly influence gene regulation. The primary focus was on two important model plant species, *A. thaliana* and *O. sativa.* The analysis was concentrated on the ∼2kb promoter regions upstream of all genes in these species, as these areas are vital regulatory elements controlling gene transcription, where cytosine methylation also plays a critical role **[29]**. Understanding the dynamics of DNA methylation in gene regulation is crucial for elucidating the molecular mechanisms that govern plant development, stress responses, and environmental adaptation.

The dataset covered 32,238 genes’ promoter sequences for *A. thaliana* and 38,309 genes’ promoter sequences for *O. sativa*. The corresponding methylation and expression data for 85 conditions (45 for *A. thaliana* and 40 for *O. sativa*) were also covered. The substantial size of these datasets (28GB) provided a strong basis for the present study and enhanced the statistical power.

Following the creation of a gene-wise dataset, the connection between gene expression and DNA methylation patterns in both species was examined by grouping genes into three distinct categories: methylated, unmethylated, and housekeeping genes. Methylated genes often showed more variable expression patterns suggesting regulatory control on them. Unmethylated genes were those for which methylation was never reported in the available data. Interestingly, unmethylated genes expressed themselves in a pattern very similar to the housekeeping genes, which are crucial for maintaining basic cellular functions. A less noisy and more sensible dataset was created by categorizing genes into these groups prior to model development. Subsequently, the expression values of all the genes in *A. thaliana* (by category) were compared and it was observed that the average expression values of methylated genes were significantly lower and highly variable than those of unmethylated and housekeeping genes (t-test *p-value*: ≤0.01), whereas the average expression values of unmethylated and housekeeping genes were nearly similar (*p-value*: 0.17) **(Fig. 3a).** The similarity between the average expression of unmethylated and housekeeping genes and their distance from that of methylated genes suggested methylation may have huge stake in defining such highly variable and spatio-temporal expression of such genes. The lesser average expression of these genes was due to their restricted expression pattern. Such genes were not expressed under most of the conditions, unlike the housekeeping and non-methylated genes. However, in certain conditions these genes expressed almost at par with the housekeeping genes (**Fig. 3a and b**), reinforcing the observation that methylation may be behind such strong spatio-temporal expression patterns. The orange violin plot shows the observed highest expression level for the methylation controlled genes, clearly suggesting that high expression is occasional and genes are under strong spatial-temporal regulation. Similar pattern was observed for both species **(Fig. 3b)**.

**Figure 3.**
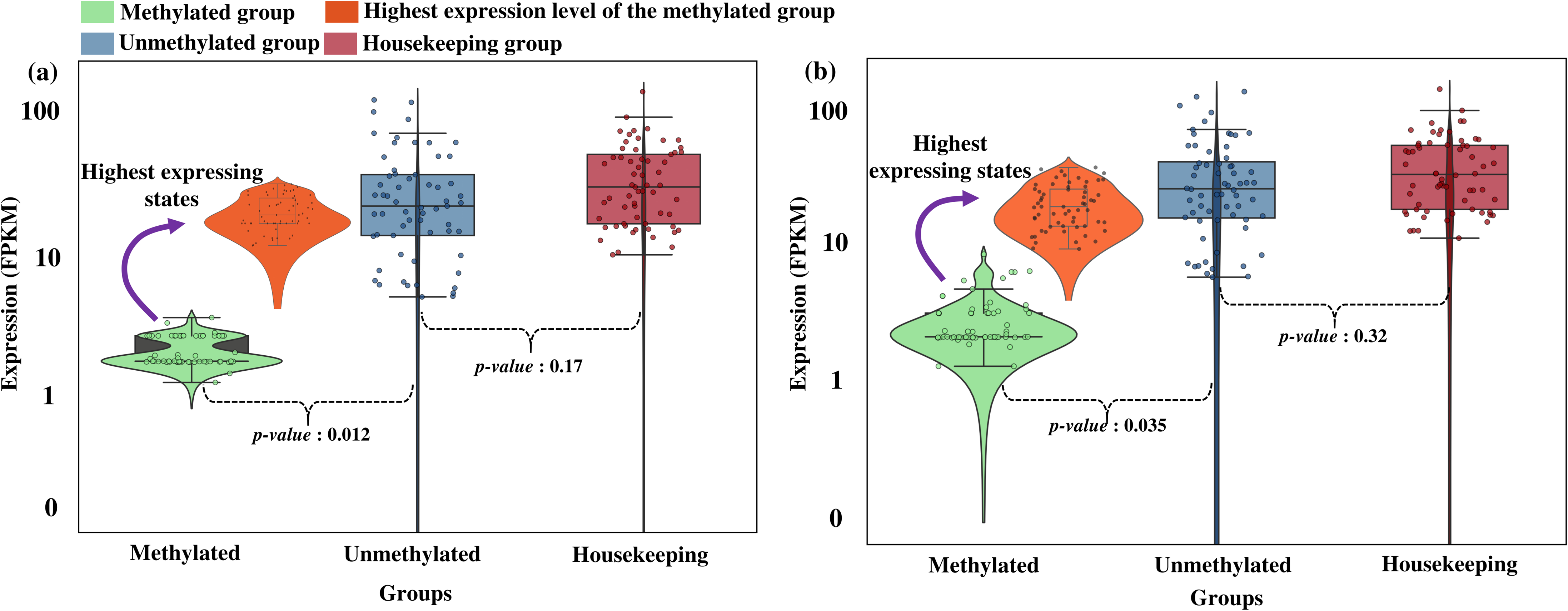
Comparative analysis of gene expression among methylated, unmethylated, and housekeeping genes. **(a)** Expression profiles in *A. thaliana* show distinct patterns across the three gene categories, with mean expression values and standard deviation error bars revealing significant differences. While unmethylated genes and housekeeping genes (internal controls) show statistically similar expression levels, methylated genes demonstrate markedly different expression patterns. The highest expression levels in methylated genes (orange violin plot) occur sporadically, indicating regulation by spatial and temporal factors. **(b)** Parallel analysis in *O. sativa* confirms conserved trends, with methylation status similarly associated with expression variation. The violin plot reveals comparable distribution patterns of highly expressed methylated genes, reinforcing that DNA methylation influences gene regulation across both model and crop species. Together, these results provide quantitative evidence for methylation-dependent expression modulation in plants, while highlighting species-specific regulatory nuances.

### Expression pattern emerges as a function of cytosine methylation pattern

The preliminary finding in the last section strongly hinted at a possible relationship between DNA methylation at cytosines and expression of the downstream genes. If any such relationship exists, patterns of methylation in the 2kb upstream may work as the domain to define a function which could produce/predict the gene’s expression level as the range. With this the problem naturally turns into a machine/deep learning problem to search such function.

The effectiveness of diverse machine learning techniques was assessed here for their aptness to relate gene expression and DNA methylation profile information. Through the assessment of multiple algorithms, including XGBoost, CNN, ResNet, and DenseNet-121, it was possible to determine the most suitable architectures through their ability to determine downstream gene expression based on methylation profile in the 2kb upstream.

The analysis began by randomly selecting 2,500 genes from both species (*A. thaliana* and *O. sativa*) and conducting training and testing on this dataset, initially with XGBoost, the highest rated shallow learning algorithm. Model parameter optimization was performed using a random grid search, resulting in the following finalized parameters: params = {“learning_rate”: 0.01, “max_depth”: 2, “subsample”: 0.1, “colsample_bytree”: 0.4, “colsample_bylevel”: 0.6, “n_estimators”: 1000}. For *A. thaliana*, the average Spearman’s correlation value of 0.94 was observed during training and 0.78 during testing **(Fig. 4a)**, where the correlation measured the relationship between the estimated and the actual experimental (observed) gene expression values. Similarly, for *O. sativa*, the average Spearman’s correlation value observed was 0.95 during training and 0.86 during testing **(Fig. 4a)**. The relationship was strong during training, though testing returned comparatively lower performance. Yet, both underlined strongly that methylation pattern in the upstream may credibly determine the downstream gene’s expression.

**Figure 4.**
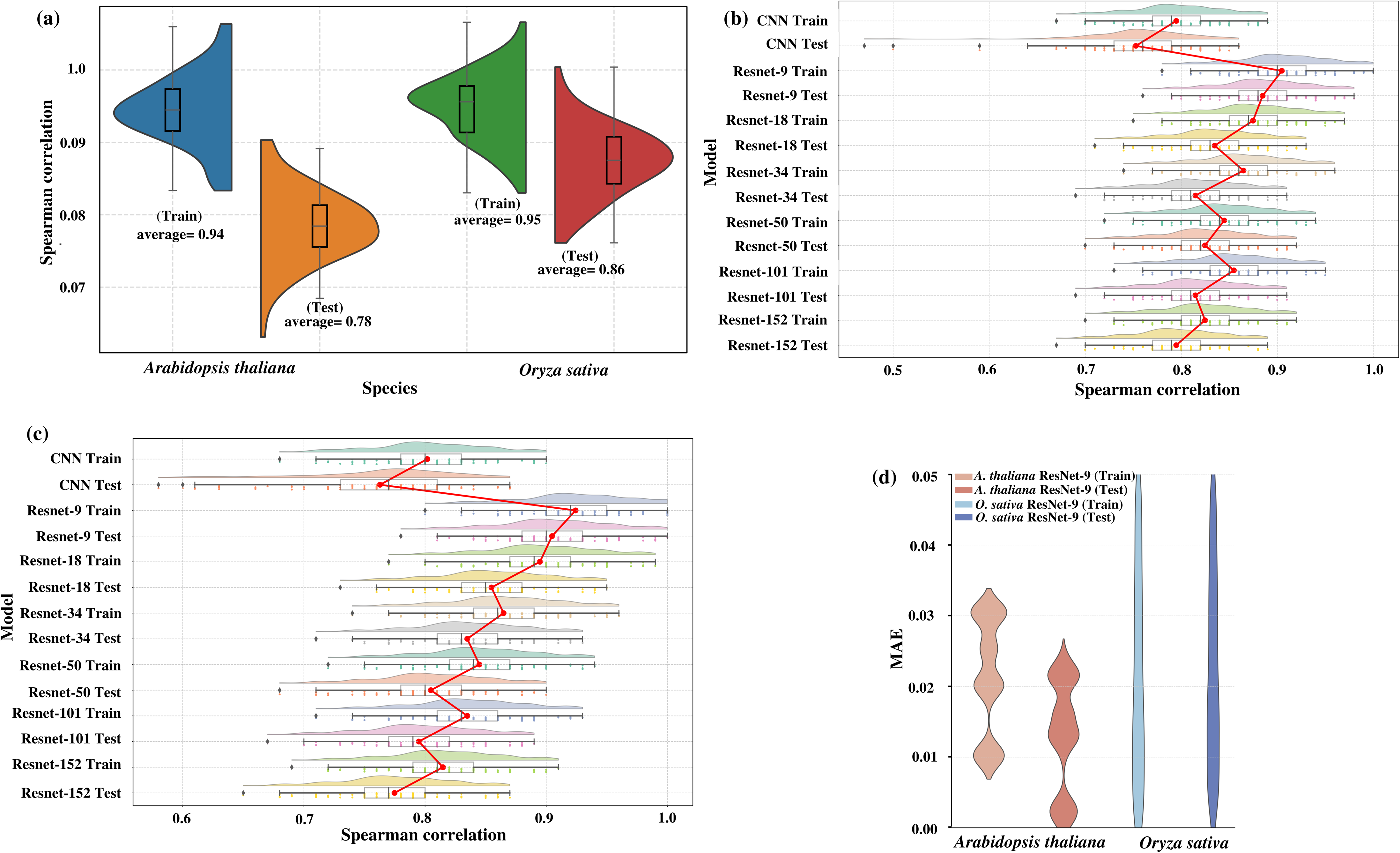
Performance evaluation of species-specific models in linking DNA methylation to gene expression. **(a)** Illustrates the model’s efficacy using XGBoost during training and testing phases. The spearman’s correlation values for both stages are presented, demonstrating the model’s capacity to elucidate the relationship between DNA methylation and gene expression. This indicates a strong correlation in the training phase, suggesting successful pattern recognition, while the testing results provide insights into the model’s suboptimal performance as significant gap in performance was noted between training and testing. **(b-c)** Comparative assessment of deep learning architectures (CNN, ResNet, DenseNet) for predicting expression from methylation patterns in *A. thaliana* and *O. sativa*, with performance quantified by Spearman’s correlation between experimental and predicted expression level patterns. These analyses identify the most effective architecture for each species. **(d)** Comparison of ResNet-9 model accuracy through Mean Absolute Error (MAE) metrics across training and test datasets, highlighting species-specific prediction capabilities.

∼10% decline in the testing performance suggested that though the machine learning was able to define a function to map methylation impact in terms of expression profile of the downstream gene, substantial overfitting was happening. This indicated the need to change the model through more efficient learning to generate better function mapping of methylation patterns to gene expression. Compared to shallow learners, deep learners can define a series of hidden functions while generating more relevant features to learn from. This makes them capable to detect features and relationships which are otherwise hard to do by shallow learners which require manual features derivation and engineering by human interventions. And latent variables cannot be appeared that way.

Subsequently, a basic Convolutional Neural Network (CNN) architecture was utilized to investigate the relationship between DNA methylation patterns and gene expression. This decision was based on the CNN’s ability to effectively extract relevant features, mitigate overfitting, accelerate training, and capability to capture spatial patterns, rendering it more suitable learning approach for exploring the complex associations between DNA methylation and gene expression. The CNN architecture is explained in the **Supplementary File Materials and Methods** section. For *A. thaliana*, the training Spearman’s correlation value was 0.79, while for the testing correlation value was 0.75 **(Fig. 4b)**. In *O. sativa*, the training Spearman’s correlation between experimental and predicted expression levels was 0.80, while the testing correlation value was 0.76 **(Fig. 4c).** With this CNN model, the training performance was poorer than the shallow XGBoost learner. Yet, the function evolved by CNN was better than the XGBoost model in a sense that it had much lesser overfitting with the performance gap between training and testing very low (∼4% only). This indicated that a deeper CNN, offering longer series of functions to functions, generating more hidden features may resolve this. But such deeper networks are also prone to information loss and vanishing gradient. This could be mitigated by using suitable architectures to counter such issues like ResNet and DenseNet, which have capabilities to carry forward the learning from previous steps much efficiently.

In the ResNet architecture, the skip connection (or residual connection) is integrated into layer, constituting a core element of the design. Skip connections allow the input of a layer to bypass one or more intermediate layers and be added directly to the output of a deeper layer. This mechanism addresses the vanishing gradient problem by enabling smoother gradient flow during backpropagation, which facilitates the training of very deep networks. By preserving the original input information and adding it to the transformed output, skip connections help the network learn residual mappings (i.e., the difference between the input and output), making it easier to optimize and improving overall performance. This design is particularly effective in capturing complex relationships like those between DNA methylation patterns and gene expression.

The same dataset was applied to the various ResNet architectures, including ResNet-9, ResNet-18, ResNet-34, ResNet-50, ResNet-101, and ResNet-152. Among these architectures, ResNet-9 exhibited the best performance. For *A. thaliana*, ResNet-9 achieved a Spearman’s correlation value of 0.90 between predicted and experimental expression level during training and 0.88 during testing **(Fig. 4b)**. For *O. sativa*, the values were 0.92 (training) and 0.90 (testing) **(Fig. 4c)**. Comparative Spearman’s correlation values for all ResNet architectures are shown in **Fig. 4b and c**, with detailed results in **Supplementary File 2, Sheet 6**. Notably, all architectures except ResNet-9 exhibited overfitting, with a 5-6% gap between the training and testing performance. Compared to the rest, ResNet-9 exhibited the least and insignificant overfitting, having MAE values of 0.0479 (training) and 0.0512 (testing) for *A. thaliana* and 0.0348 (training) and 0.0382 (testing) for *O. sativa* **(Fig. 4d)**. The small differences between train and test performance (0.0033 for *A. thaliana* and 0.0034 for *O. sativa*) with statistically insignificant p-values (p = 0.1362 for *A. thaliana* and p = 0.1284 for *O. sativa*) aptly underlined the robustness and generalization capability of the raised ResNet-9 based model.

To further validate these findings, 10-fold random trials of training-testing was performed on randomly selected 10 genes from both species. The results showed consistent MAE values across the folds, and a t-test comparing the training and testing MAE values yielded insignificant p-values (p > 0.05), reinforcing the robustness of the model and the absence of any significant overfitting. This investigation elucidates the superior capacity of the developed deep-learning model to map the underlying relationship between DNA methylation and gene expression, suggesting that deeper architectures are more efficacious in establishing these relationships. The details of the implemented deep-learning architecture are given in the methods section and **Fig. 2b**.

After conducting hyperparameter optimization, the model performance for both species was markedly improved. Prior to optimization, the average Spearman’s correlation for testing in *A. thaliana* was 0.88. Post-optimization, this value increased to 0.94. Similarly, for *O. sativa*, the Spearman’s correlation for testing improved from 0.90 to 0.95 **(Fig. 5a)**.

**Figure 5.**
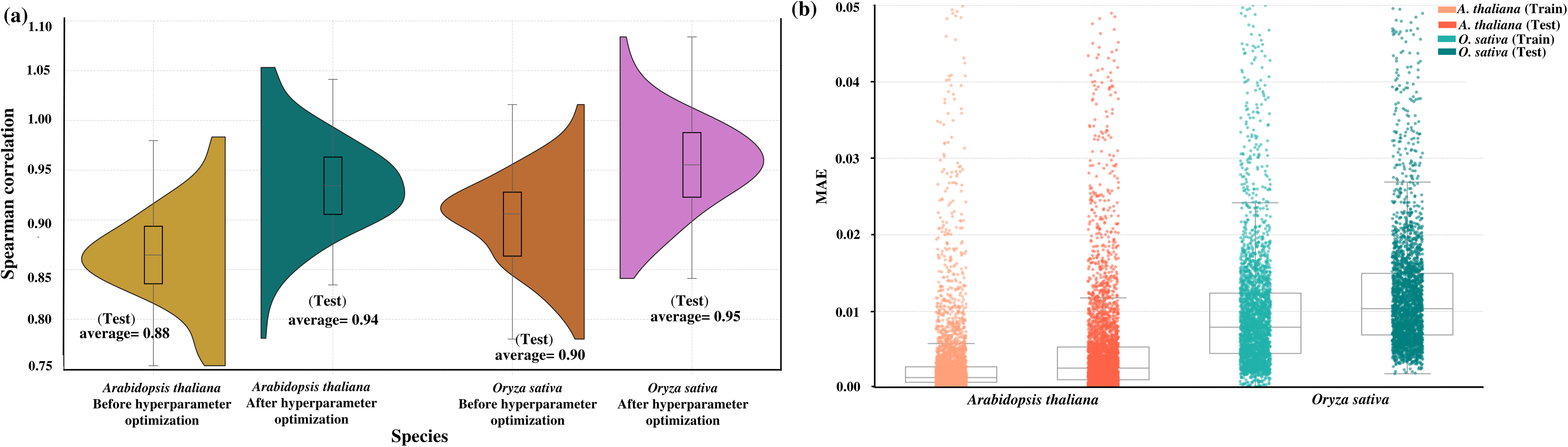
Hyperparameter optimization and model performance analysis. **(a)** Comparison of model testing performance before and after hyperparameter optimization for both A. thaliana and *O. sativa.* **(b)** Quantitative assessment of optimization effects shown through bar graphs of either Spearman’s correlation coefficients (ρ) or Mean Absolute Error (MAE) values. The significant performance improvement following optimization demonstrates the critical role of parameter tuning in enhancing both predictive accuracy and model generalizability for studying methylation-expression relationships across species.

For all the modeled genes in *A. thaliana*, the developed deep-learning model achieved average MAE values of 0.0389 during training and 0.0406 during testing. Likewise, for *O. sativa*, the model demonstrated average MAE values of 0.0312 in training and 0.0354 in testing **(Fig. 5b)**. A t-test comparing the training and testing MAE values yielded insignificant p-values, further supporting the absence of any significant differences between the performance on training and testing datasets. With this all, it was now clearly established that upstream cytosine methylation pattern guide a gene’s expression levels. And this could be precisely learned by deep-learning, which in turn was able to accurately estimate the gene expression levels under variable conditions with variable methylation patterns.

### Development of a universal model: GC content similarity holds the key

By now, this was clear that deep-learning was highly successful in establishing the relationship between variability in cytosine methylation pattern in the upstream region to the genes and their expression levels. However, plants exhibit significant diversity across species, particularly in transcriptional regulation, as highlighted in recent studies **[30–32]**. This diversity suggests that a model trained for a gene in one species may not perform with the same efficacy when applied to another species. To address this, the universal capability of the developed deep-learning model was assessed by focusing on homologous genes across species. The initial approach involved using homology to pick the suitable ones from one species and predict the corresponding gene expression in other species. It was hypothesized that homologous genes would retain same functional characteristics, and thus should express in similar fashion with similar regulatory control.

Initially, five random gene models were selected from *A. thaliana* which were tested across their homologous counterparts in *O. sativa*, and vice versa. Within species testing yielded strong average Spearman’s correlation coefficients between predicted and actual expression levels (0.82 for *A. thaliana* **(Fig. 6a)** and 0.81 for *O. sativa* **(Fig. 6b))**. However, cross-species testing revealed a sharp decline in the average correlation values: 0.61 for *A. thaliana* models applied to *O. sativa* **(Fig. 6a)** and 0.60 for *O. sativa* models applied to *A. thaliana* **(Fig. 6b)**. This indicated that gene homology does not guarantee similar transcriptional regulation. Homologous genes appeared to have different regulation controls. To investigate further, the 2kb upstream regions of these homologous genes were compared. Their GC% differed significantly. Since DNA methylation in plants is cytosine-focused, it was hypothesized that GC% might play a critical role in gene regulation. Thus, focus was shifted towards grouping the genes with similar GC% in their upstream regions, aiming to develop models based on GC content similarity instead of homology.

**Figure 6.**
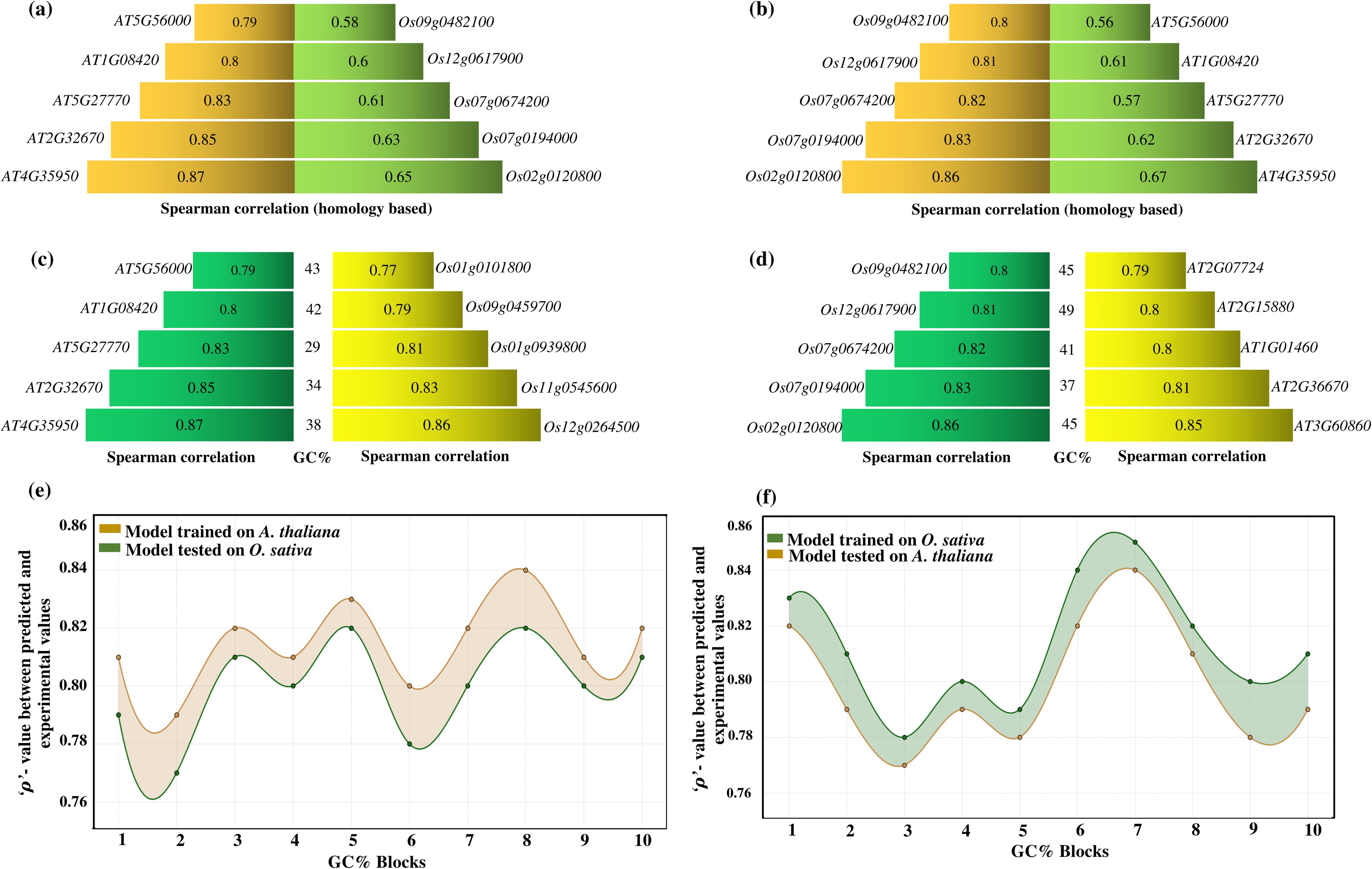
Construction of a universal model. **(a)** Performance evaluation of five *A. thaliana* genes and their *O. sativa* homologs shows a strong Spearman’s correlation (average ρ = 0.81) between predicted and experimental expression values within species but a significantly reduced cross-species performance (average Δρ = 0.60), indicating limited conservation of regulatory mechanisms. **(b)** Reciprocal analysis of five *O. sativa* genes and their *A. thaliana* homologs confirms this species-specificity pattern. **(c-d)** GC%-stratified analysis reveals that models trained on genes with similar GC content maintain high predictive accuracy (average ρ = 0.82) both within and across species, suggesting GC% as a key determinant of regulatory conservation. **(e-f)** Group-level analysis demonstrates that gene blocks with similar GC% exhibit exceptional cross-species performance (average ρ = 0.81), supporting the effectiveness of GC%-guided universal modeling.

To systematically assess the impact of GC% across species, the GC% of the 2kb upstream regions for the five randomly selected genes in *A. thaliana* were calculated. Following this, five genes in *O. sativa* with similar GC% to those in *A. thaliana* were selected. Separate predictive models were trained for each *A. thaliana* gene, and their performance was evaluated within the species. The results changed drastically with a strong average Spearman’s correlation coefficient of 0.82 between the predicted and actual expression levels **(Fig. 6c)**, demonstrating the effective predictive power within species even if they differed in homology but had closer GC% in their promoters. Interestingly, when these trained models were applied to the *O. sativa* genes with closest GC content, the correlation remained high at 0.81 **(Fig. 6c)**, closely resembling the performance observed within *A. thaliana*. A similar trend was observed when *O. sativa* models were tested on *A. thaliana* genes with similar GC% **(Fig. 6d)**. This consistency suggested that GC content is a highly significant factor for predicting relationships between DNA methylation pattern and gene expression across different plant species.

After observing these promising results for the five genes based on GC%, genes with similar GC% value ranges were grouped into blocks. This approach was adopted because building individual models for each gene with a similar GC% would be highly time consuming and computationally expensive process. Grouping genes with a similar GC% allowed development of generalized models which could predict the expression of multiple genes within the same GC% range group, leveraging from shared regulatory patterns. To validate this approach, the grouped models of *A. thaliana* and *O. sativa* were tested. Models trained on *A. thaliana* and tested on *O. sativa* achieved an average Spearman’s correlation of 0.81 between actual and predicted expression levels **(Fig. 6e)**, while models trained on *O. sativa* and tested on *A. thaliana* achieved an average correlation of 0.80 **(Fig. 6f)**. These results demonstrate that grouping genes with similar GC% significantly improved the cross-species predictive performance, underscoring the importance of GC% in establishing relationships between DNA methylation pattern and gene expression.

The findings highlight that promoter region GC content, not homology, is a key determinant of regulatory behavior and gene expression. This suggests that GC content-based models can significantly enhance cross-species predictive accuracy. While homologous gene sequences provide insights into functional conservation within the genic regions, they fail to account for regulatory similarities that GC content were found capable to explain.

### Reversing the wheel: Using expression to determine the critical cytosines

Upto here, this was established that upstream cytosine methylation can effectively predict a gene’s expression, and deep-learning model discussed above achieved that. Identifying the specific cytosines essential for gene regulation can provide valuable insights into a gene’s regulatory switches and reveal potential targets among them to manipulate a gene’s expression. Cytosine methylation is dynamic. Its status can change under various environmental or biological conditions, influencing gene expression. This variability makes it challenging to identify which cytosines are critical and how they impact expression. In terms of translation, it becomes more important question than predicting expression of a gene. RNA-seq and transcriptome profiling experiments are easily approachable now and routine practices. If they could be used to identify critical cytosines influencing a gene’s expression, a bigger purpose would be achieved. Recently, we demonstrated successfully that transcriptome profiles can be used reveal the condition specific cytosines methylation states **[11]**. Thus, attempting to decode criticality of such cytosines appeared promising and a natural next step.

Two different approaches were taken: (A) in-silico deletion of cytosines and measuring its impact on predicted expression accuracy. (B) Identifying most critical cytosines through machine weights and explainable AI. In the first one, three complementary methods were applied to study the promoters and their associated gene’s expression in *A. thaliana* and *O. sativa*. Two representative genes were randomly selected for the initial analysis: UMAMIT28 (AT1G01070) from *A. thaliana* and OsHXK8 (Os01g0190400) from *O. sativa*. In the first approach, individual cytosines were knocked out from the promoter regions for their data input to the DL-model. Thereafter, such knocking out was evaluated for their impact on the models’ ability to estimate the downstream gene’s expression.

To achieve this, a baseline (actual) correlation value was first established by training the model on the promoter sequence (with all cytosines intact) and measuring its Spearman’s correlation between the actual and estimated expression levels of the downstream gene. Next, individual cytosines were knocked out and models’ performance was evaluated while measuring the Spearman’s correlation between the new estimated expression values and the actual values. A significant drop in correlation was observed after the knocking out of specific cytosines which indicated that these positions played a critical regulatory role. Cytosines whose knocking out had minimal impact on correlation were deemed less influential in controlling the gene’s expression. Ranking cytosines based on their impact allowed identification of key regulatory sites. For instance, in *A. thaliana* (AT1G01070 or UMAMIT28), knocking out cytosines at positions 1,787 and 1,869 reduced the Spearman’s correlation from 0.72 to 0.58 and 0.72 to 0.56, respectively, indicating their potentially strong regulatory role. In contrast, knocked out cytosines at positions 1,041 and 1,022 led to a smaller decline, from 0.72 to 0.69 and 0.72 to 0.70, respectively **(Fig. 7a)**. The dotted red line in **Fig. 7a and b** represents the baseline Spearman’s correlation before any knockout. A similar trend was observed for *O. sativa* (Os01g0190400 or OsHXK8), as shown in **Fig. 7b**, These findings emphasize that not all cytosines contribute equally to gene regulation and only a subset of cytosines serves as critical regulatory points. Identifying these key cytosines sites provides an important gateway towards better understanding of epigenetic regulatory switches, which can be effectively used for human interventions and purposes.

**Figure 7.**
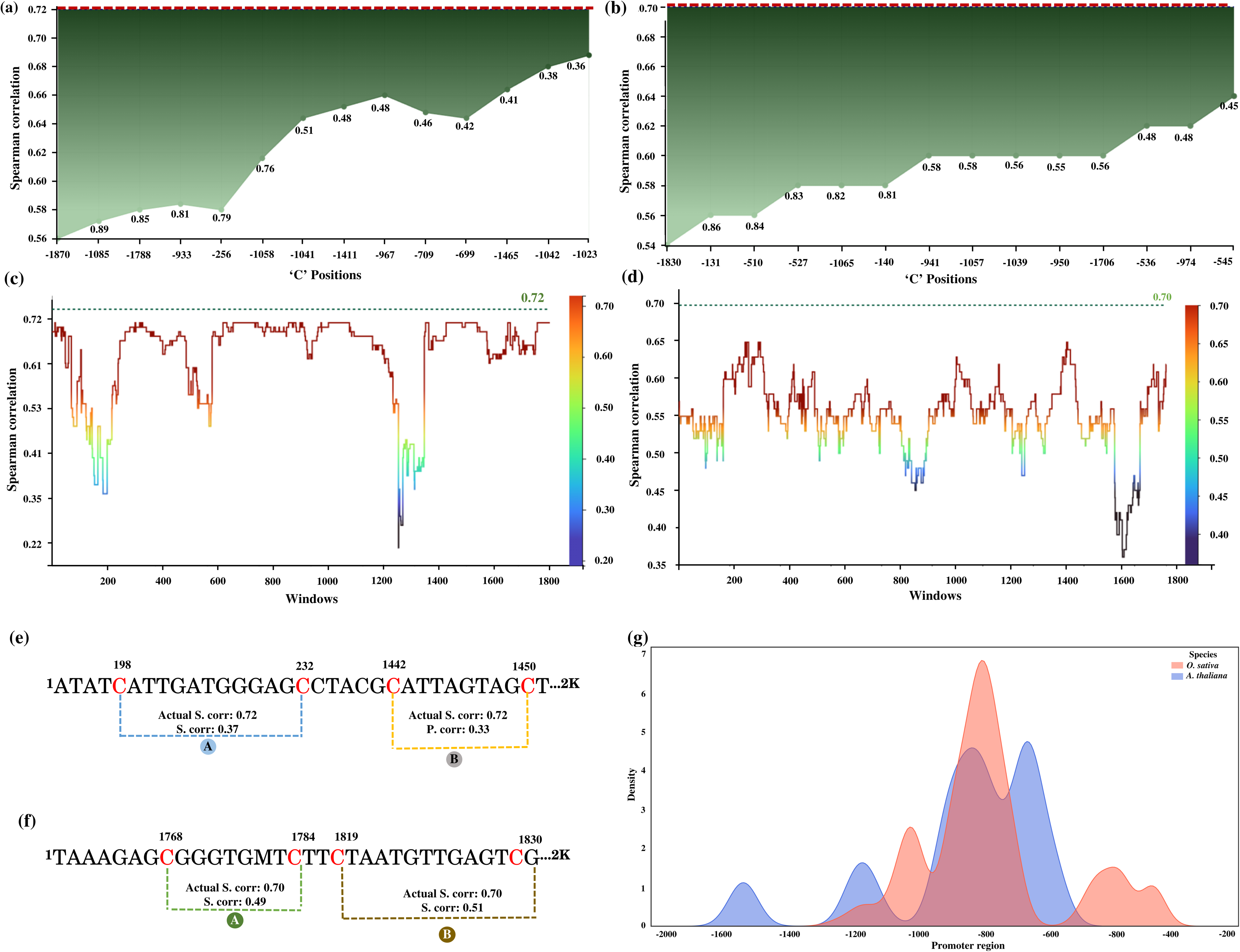
Identification of critical cytosines and their distribution patterns in the promoter region. **(a-b)** The impact of systematically knocking out individual cytosines in the promoter region of *A. thaliana* (AT1G01070 or UMAMIT28) and *O. sativa* (Os01g0190400 or OsHXK8) on gene expression. The graph shows a reduction in Spearman’s correlation, with the actual correlation indicated by a red dotted line. The associated Grad-CAM score at the knocked-out cytosine position (black digits) highlights the cytosine’s importance in gene expression regulation, emphasizing the model’s focus on this site. **(c-d)** A 100 bp overlapping window approach was applied to evaluate how groups of cytosines collectively influence gene regulation in both species. In *A. thaliana* (AT1G01070 or OsHXK8), cytosines within window numbers 190–205 (corresponding to base positions -1810 to -1795 relative to the TSS) had a substantial impact on gene expression, as Spearman’s correlation dropped sharply from 0.72 to 0.33 upon knockout. A similar effect was observed in window 1,372–1,373 (base positions -628 to -627 from the TSS), indicating that specific promoter regions contain cytosines crucial for regulation. A comparable trend was observed in *O. sativa* (Os01g0190400 or OsHXK8). **(e-f)** A cytosine combination approach was used to examine how cytosine groupings influence gene expression. In *A. thaliana*, combination A (knocking out cytosines at positions 198 and 232) reduced Spearman’s correlation from 0.72 to 0.37. Similarly, the impacts were seen for the other given combinations including rice. **(g)** KDE plots illustrating the positional preferences for critical cytosines.

In the next analysis, the 100 bp overlapping windows were evaluated across the promoter regions to assess if such critical cytosines might exist in continuous groups too. This helped in knowing the region specific criticality. **Fig. 7c** illustrates the results for *A. thaliana* (AT1G01070 or UMAMIT28), where the X-axis represents the window numbers, and the Y-axis displays the Spearman’s correlation values between actual and predicted gene expression. Notably, cytosines within window numbers 190-205 (corresponding to base positions -1810 to -1795 relative to the TSS) had a substantial impact on gene expression, as the Spearman’s correlation dropped sharply from 0.72 to 0.33 upon being knocked out. A similar effect was observed in window numbers 1,372–1,376 (base positions -628 to -624 from the TSS), further indicating that specific promoter regions harbor cytosines crucial for regulation. A comparable trend was observed in *O. sativa* (Os01g0190400 OsHXK8), as shown in **Fig. 7d**. By analyzing the combined influence of cytosines within each window, it was also clear that their are regions in promoters which are enriched for critical cytosine in form of blocks.

The third and final approach focused on identifying cytosine pairs and their combinations to examine how these groupings influenced gene expression. We particularly focused on the pairs which when knocked out, led to a significant decrease in expression while alone they didn’t appeared critical. As shown in **Fig. 7e** for *A. thaliana*, combination “A” knocking out cytosine at positions 198 and 232 together, resulted in a Spearman’s correlation reduction from 0.72 to 0.37. Similarly, in combination “B”, knocking out cytosine at positions 1442 and 1450 decreased the correlation from 0.72 to 0.33. In **Fig. 7f** for *O. sativa*, the Spearman’s correlation for combination “A” decreased from 0.70 to 0.49, while for combination “B”, it got reduced to 0.51. This highlights the importance of cytosine combinations in gene regulation, where the effect of one cytosine may be dependant upon the presence of nearby cytosines. Individually, these cytosines were not scoring high on influence. But when paired with other cytosines, their impact becomes visible. This all clearly suggest qualitative as well as quantitative impact of cytosines on gene expression.

Critical cytosines were identified through the systematic knockout of cytosines, both individually and in groups. Systematic knocking out of cytosines and their groups is helpful in annotating the existing genes and developing knowledge about them, storing that in term of databases. However, it is a time consuming task and not pragmatic for on the spot detection of criticality. To overcome these limitations, a second way was employed my two approaches: (I) Extraction of importance weights from the trained model, (II) Explainability AI based approach, Grad-CAM.

The first method involved employing in-house scripts to extract cytosine weights from the eighth convolutional layer of each gene model. These weights were compared with the corresponding Spearman’s correlation values to determine whether cytosines deemed critical exhibited higher weights. For the second approach, Grad-CAM score profiles were implemented using the eighth convolutional layer of the deep-learning system architecture as the focus to identify the most significant cytosines influencing gene expression. This layer is crucial because it captures the most significant features (cytosines) affecting gene expression. By concentrating on this layer, the critical cytosines which play a key role in regulating gene expression could be effectively identified. This analysis revealed a strong correlation between Grad-CAM scores and Spearman’s correlation values, confirming that cytosines identified as critical ones exhibited the largest drop in Spearman’s correlation upon knockout while also having high Grad-CAM scores, as illustrated in **Fig. 7a and 7b**. This way the Grad-CAM approach was ultimately implemented for the automated, real-time detection of critical cytosines. While the knockout methods were conducted on AT1G01070 and Os01g0190400, we validated these observations across multiple genes in both species, confirming that critical cytosines are gene and position specific and that not all cytosines contribute equally to gene regulation. All gene-specific, information for both species has been systematically stored in a database, discussed in an upcoming section below.

Following the identification of gene-specific critical cytosines in both *A. thaliana* and *O. sativa*, their distribution was examined within promoter regions to determine whether these critical cytosines preferred any specific genomic regions when analyzed collectively. Analysis revealed that though the critical cytosines destributions differed in both species, a significant amount of critical cytosines were concentrated within the 1kb upstream of the transcription start site (TSS). This region is well known for its importance in gene expression regulation **(Fig. 7g)**. This observation aligns with previous studies indicating that gene regulatory elements, including promoters, enhancers, and transcription factor binding sites, are typically enriched within this proximal upstream region **[33]**.

### Critical cytosines mirror drift in functional priority across species

As transpires now, there are critical cytosines which influence downstream gene’s expression. They work either quantitatively or qualitatively, and display certain positional bias also. The genes across species are functionally resolved on the basis of homology. However, the present study found that in terms of regulation, it is GC% similarity in the promoter regions, which explains their regulation and expression.

To further understand the functional significance of genes containing the critical cytosines, their Gene Ontology (GO) annotations and functional associations, as well as their functional dynamics shifts between the two species were analyzed. Comparing GO results between *A. thaliana* and *O. sativa* could provide insights into conserved regulatory mechanisms or species-specific adaptations in gene regulation mediated by these critical cytosines.

In this study, all genes containing critical cytosines in both species were ranked based on the presence of these cytosines in their promoter regions, along with their expression Spearman’s correlation and Grad-CAM scores. From this ranking, the top 400 genes were selected in each species, prioritizing those where critical cytosines had the most significant regulatory impact. These selected genes, highly enriched with critical cytosines, served as the ideal candidates for understanding the regulatory drifts. GO enrichment and pathway analysis were performed, identifying key biological processes, cellular components, and molecular functions associated with these genes.

The Gene Ontology (GO) analysis of the genes enriched for critical cytosines in *A. thaliana* and *O. sativa* highlights distinct yet interconnected regulatory roles of critical cytosines in shaping plant development and metabolic functions. In *A. thaliana*, these genes are predominantly linked to ribosomal DNA (rDNA) maintenance, cellular development, and stamen development, suggesting a strong influence on genome stability and reproductive success. rDNA plays a crucial role in nucleolar integrity by serving as the scaffold for nucleolus formation **[35,35].** It defines the last leg of gene expression through mRNA translation. Stamen development, governed by cytosine methylation, directly influences pollen viability and fertility, supporting the role of epigenetic modifications in and control of a time dependent event of reproduction. A comprehensive breakdown of these results can be found in **Table S2 of Supplementary File 1**. Molecular function and pathways associations supported this observations, where enrichment for phytohormone modulations, histone modifications, and primary metabolic processes was found. Localization of these genes to the endopeptidase complex, early endosomes, and generative cell nuclei further indicates their involvement in protein degradation, intracellular trafficking, and reproductive cell differentiation all associated with the process of regeneration. In overall all these enriched genes are highly spatio-temporal and orchestral in their fundamental roles. Having critical cytosines enrichment further justifies the high stakes of epigenetic control on them.

In contrast, *O. sativa* exhibits a markedly different pattern, with genes enriched for critical cytosines predominantly associated with phospholipid biosynthesis, isopentenyl diphosphate biosynthesis, and carbon fixation, reflecting a strong emphasis on metabolic regulation and photosynthetic efficiency, storage and energy management. The enrichment of shikimate kinase activity in *O. sativa* further reinforced the role of critical cytosines in secondary metabolism, particularly in the synthesis of aromatic amino acids, which serve as precursors for a wide range of secondary metabolites essential for plant defense, signaling, and growth. A more detailed description of these findings is presented in **Table S3 of Supplementary File 1**.

A comparative analysis of both species reveals key distinctions in the regulatory functions influenced by critical cytosines. While *A. thaliana* prioritizes developmental regulation, particularly in reproductive and cellular differentiation processes, *O. sativa* exhibits a strong metabolic focus, emphasizing energy production, lipid biosynthesis, and photosynthetic efficiency. These differences likely reflect evolutionary adaptations tailored to each species’ unique growth environments and physiological demands.

To investigate the conservation of essential gene functions between *A. thaliana* and *O. sativa*, a comparative analysis of the genes enriched for critical cytosines in both species was conducted with broader grouping. This study was expanded to cover the top 1,500 critical cytosine enriched genes. These 1,500 genes were categorized into three equal groups of 500 genes (high, medium, and low criticality) to systematically examine the distribution of functional associations of these three levels of criticality.

In terms of biological processes, distinct patterns of gene enrichment were observed in *A. thaliana* and *O. sativa*, reflecting their unique evolutionary and ecological adaptations. In the high criticality group, 12 genes in *A. thaliana* were highly enriched in the maintenance of the rDNA process, which is a critical function for translation and ribosomal biogenesis **(Fig. 8a)**. This is particularly important in *A. thaliana*, a model plant with a compact genome and rapid life cycle, in which efficient ribosome production is essential for growth and development. In contrast, in *O. sativa*, 16 genes enriched in the same process dropped down into the medium criticality group, suggesting that rDNA maintenance, although still important, may be less prioritized in rice because of its larger genome size, polyploidy, and reproductive strategy focused on seed production. A total of 36 genes in *A. thaliana* and 22 genes in *O. sativa* from the high criticality group were enriched in cellular development process. This indicates that these genes play a conserved role across both species, but the extent of their criticality differs. Similarly, in the medium-criticality group, 19 genes in *A. thaliana* and 32 genes in *O. sativa* (which were classified as high-criticality) were enriched in cell differentiation processes. This observation suggests that while both species share this biological function, the regulatory importance of critical cytosines differs between them. The greater number of *O. sativa* genes in the high-criticality group suggests that critical cytosines may exert a stronger regulatory influence on cell differentiation in this species compared to *A. thaliana*. Biological processes such as cellular metabolism, transferase activity, and pyruvate metabolic processes were enriched in the low criticality group for both species, indicating that these fundamental processes are less influenced by critical cytosines, highlighting their ominipresent roles in basal activities where spatio-temporal regulation is not much required. Regarding cellular components **(Fig. 8b)**, the endopeptidase complex was associated with the low criticality group in *A. thaliana*, but with the high criticality group in *O. sativa*, suggesting species-specific roles in protein degradation and regulation. In rice, the endopeptidase complex may play a critical role in stress responses, such as drought or pathogen resistance, which are essential for survival under diverse agroecological conditions.

**Figure 8.**
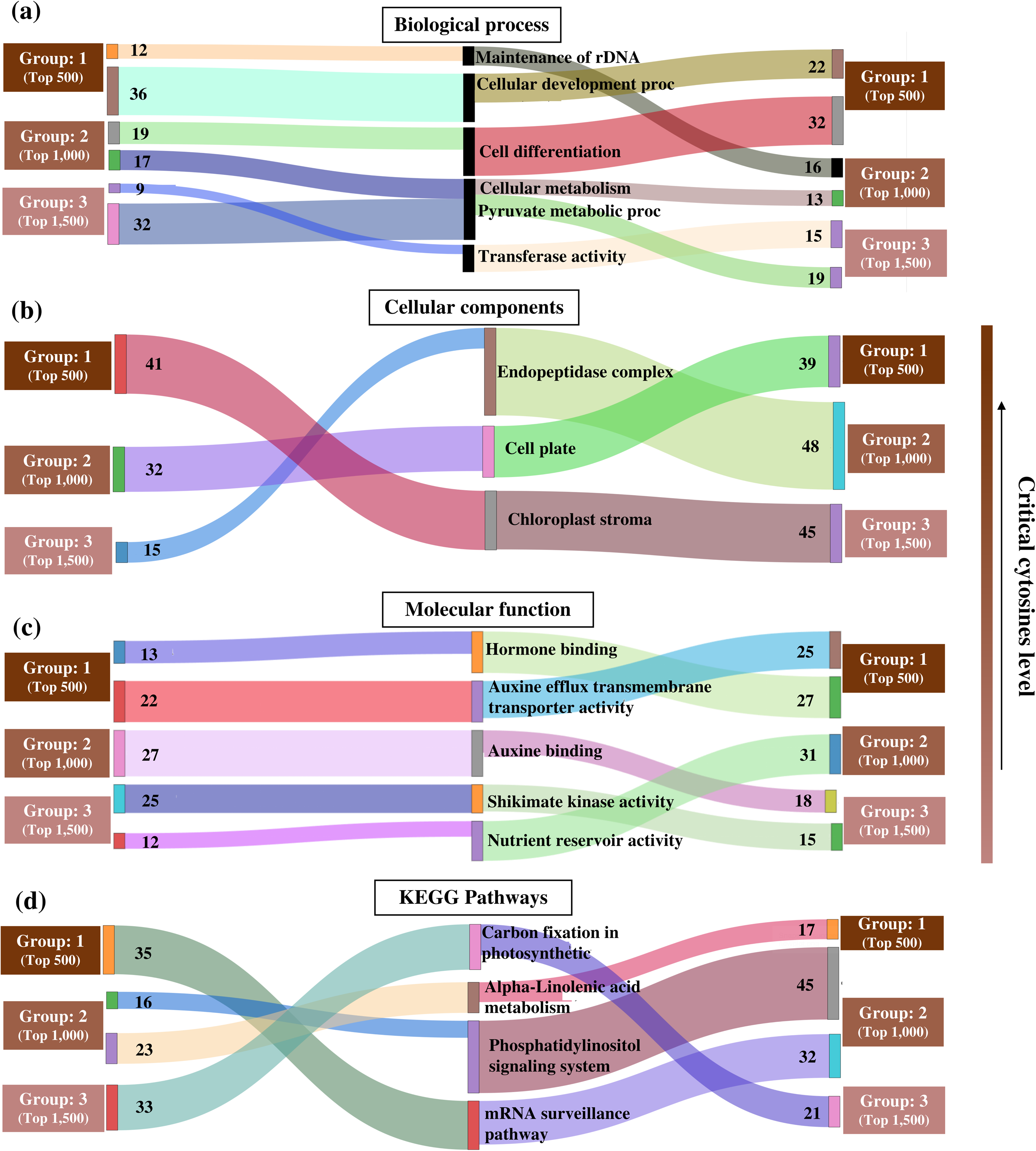
Sankey plot illustrating critical cytosines mirror drift in functional priority across species. The diagram highlights conserved and divergent gene functions, demonstrating how shifts in criticality reflect species-specific adaptations to ecological and evolutionary pressures. These insights underscore the role of genes containing critical cytosine and epigenetic regulation in plant adaptation, **(a)** Biological Processes: rDNA maintenance is highly critical in *A. thaliana* but moderately critical in *O. sativa*, reflecting differences in genome organization and reproductive strategies. **(b)** Cellular Components: Chloroplast stroma is highly critical in *A. thaliana* but less so in *O. sativa*, likely due to rice’s adaptation to high-light environments and its emphasis on starch biosynthesis, **(c)** Molecular Functions: Hormone binding and auxin transport are highly critical in both species, whereas nutrient reservoir activity is more critical in *O. sativa*, aligning with its role as a cereal crop, **(d)** KEGG Pathways: mRNA surveillance is highly critical in *A. thaliana* but moderately critical in *O. sativa*, suggesting species-specific post-transcriptional regulation. Alpha-linolenic acid metabolism is more critical in *O. sativa*, likely due to its reliance on lipid metabolism for stress responses.

In the molecular function category **(Fig. 8c)**, hormone-binding and auxin efflux transmembrane transporter activity were enriched in the high criticality group for both species, underscoring the conserved importance of hormone signaling in plant growth and development, which is universally spatio-temporal and needs to be tightly regulated. In contrast, nutrient reservoir activity was associated with the low criticality group in *A. thaliana*. While it was in the medium criticality group in *O. sativa*, reflecting rice’s adaptation as a cereal crop, with a focus on nutrient storage in seeds. This all difference highlight the regulatory drifting between a model and a crop species, where nutrient storage is a key trait for agricultural productivity.

The mRNA surveillance pathway was enriched in the high criticality group for *A. thaliana* **(Fig. 8d)** which belonged to the medium criticality group for *O. sativa*, indicating differences in post-transcriptional regulation between the two species. In *A. thaliana*, precise control of mRNA stability and translation may be critical because of its rapid life cycle and the need for efficient resource allocation. This was paralleled by rRNA genes enrichment in *A. thaliana,* as observed above. Likewise, alpha-linolenic acid metabolism was associated with the medium criticality group in *A. thaliana*, but moved into the high criticality group in *O. sativa*, most likely due to rice’s reliance on lipid metabolism for energy storage. Alpha-linolenic acid is also a precursor for jasmonic acid, a hormone involved in defense responses.

Overall, this study enhanced our understanding of the role of critical ‘C’ in imparting species specific modulation of gene expression and its control. Observations made here pave the way for future studies aimed at unraveling the molecular underpinnings of plant adaptation and resilience, where cytosines and epigenetic control now appear as highly imminent regulators.

### Cross-Species validation extends model applicability to diverse plant families

To further validate the developed algorithm’s (CritiCal-C) universal applicability beyond *A. thaliana and O. sativa*, the model was applied to three agronomically important species representing diverse plant families: *Brassica rapa* (*Brassicaceae*), *Cucumis sativus* (*Cucurbitaceae*), and *Solanum tuberosum* (*Solanaceae*). For each species, 2 kb promoter sequences were extracted genome-wide, methylated cytosines were marked based on publicly available whole-genome bisulfite sequencing data, and CritiCal-C was used to identify critical cytosines and predict gene expression without species-specific retraining. The top 1,500 genes ranked by critical cytosine enrichment (based on Grad-CAM scores) were subjected to Gene Ontology enrichment analysis to assess functional conservation and species-specific adaptations.

The analysis revealed both conserved regulatory themes and species-specific functional priorities across the five plant species examined. All species showed enrichment of critical cytosines in genes associated with hormone signaling and cellular development, confirming universal regulatory importance of these processes **(Supplementary File Materials and Methods, Fig. S4)**. However, species-specific patterns emerged that reflected distinct ecological and agronomic adaptations: *B. rapa* exhibited enrichment in glucosinolate biosynthesis and defense responses, consistent with its Brassicaceae-specific secondary metabolism; *C. sativus* showed strong enrichment in fruit development and water transport processes, aligning with its cucurbit-specific developmental programs; and *S. tuberosum* displayed enrichment in starch biosynthesis and tuber development, reflecting its adaptation for underground storage organ formation **(Supplementary File 2, Sheet 7-9, Supplementary File Materials and Methods, Fig. S5 and S6)**. These findings demonstrate that while CritiCal-C successfully identifies functionally critical cytosines across phylogenetically distant species without retraining, the specific biological processes under critical cytosine control reflect species-specific evolutionary adaptations, validating the model’s capacity to capture both universal regulatory principles and lineage-specific epigenetic control mechanisms.

### Experimental validation of critical cytosine identification

AT1G20400, encoding a DUF1204 domain-containing protein involved in cellular organization in *A. thaliana*, was selected for experimental validation. CritiCal-C identified seven cytosines within a 100 bp promoter window (positions 794-894 bp upstream of the transcription start site) showing potential regulatory cytosines with high criticality scores.

Quantitative RT-PCR revealed strong downregulation of AT1G20400 under heat stress compared to 22°C controls (**Supplementary File Materials and Methods, Fig. S7**). Expression dropped 5.8-fold (log₂ fold-change = -2.5, p < 0.01, Student’s t-test), ranking among the most heat-responsive genes tested alongside SESA4, NAP4, QHNDH, BRD5, SESA2, LTP1, and Hexokinase **(Fig. 9a)**. This pronounced transcriptional response suggested active regulatory remodeling during heat stress, making AT1G20400 suitable for testing whether CritiCal-C correctly identified functionally important cytosines.

**Figure 9.**
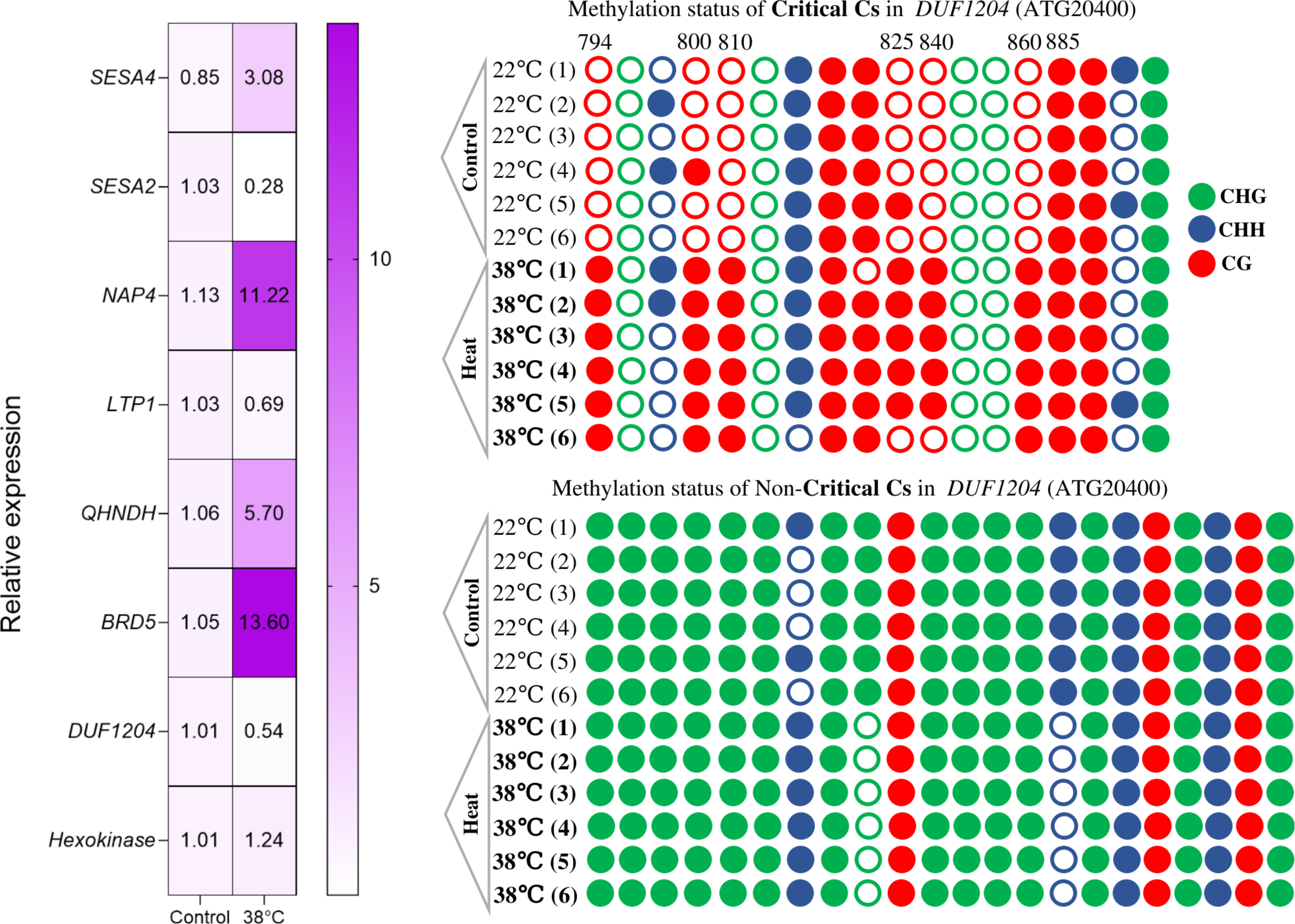
Experimental validation of computationally identified critical cytosines in DUF1204 under heat stress **(a)** Expression profiling of the genes under heat stress using qRT-PCR. **(b)** and **(c)** Effect of heat stress on the methylation of cytocines in Critical **(b)** and non-critical **(c)** 100bp window. Filled circles correspond to methylated CpGs; unfilled circles correspond to unmethylated CpGs.

Targeted bisulfite sequencing of the critical window (794-894 bp) revealed striking methylation patterns **(Fig. 9b)**. Six of the seven identified critical cytosines underwent dramatic methylation increases under heat stress. At 22°C, these six positions showed minimal baseline methylation, only 1-2 of 6 sequenced clones carried methylation. Heat stress triggered extensive hypermethylation, with 5-6 of 6 clones methylated at these sites. All six responsive cytosines occurred in CG dinucleotide contexts, the most stable methylation form in plants. Primer sequences used for bisulfite PCR amplification and qRT-PCR analysis are provided in **Supplementary File 2, Sheet 9.**

CritiCal-C’s knockout analysis assigns criticality based on performance impact: cytosines causing substantial Spearman correlation drops when knockout play critical regulatory roles, while those causing minor drops do not. The model generates post-knockout correlation values, where lower value indicate higher criticality. In single-cytosine knockout tests, six positions showed low post-knockout correlations-C794 (0.30), C800 (0.38), C810 (0.31), C825 (0.39), C840 (0.38), and C860 (0.33) predicting high methylation responsiveness. Experimental validation confirmed this identifications: these six positions underwent hypermethylation under heat stress. The seventh cytosine, C885 (0.43), despite residing in the same promoter region, showed no methylation change between control and stress conditions, consistent with its borderline criticality score.

Pairwise knockout analysis reinforced these findings. C885 exhibited a slight correlation drop from the baseline value when knockout with neighboring cytosines, whereas the six responsive positions showed substantial correlation drops when paired with neighbors. The C794-C800 and C825-C840 pairs demonstrated particularly strong synergistic effects, with combined knockouts producing greater impacts than individual removals. This hierarchical pattern, where lower post-knockout correlations corresponded to stronger methylation responses demonstrates that CritiCal-C’s numerical rankings capture genuine regulatory differences rather than arbitrary classifications.

To verify specificity, the non-critical control window (positions 0-100 bp) within the same AT1G20400 promoter underwent bisulfite sequencing. This region contained cytosines with high post-knockout correlations (>0.70), indicating minimal identified regulatory importance. Among 22 cytosines spanning this window, 18 showed no methylation differences between control and heat stress conditions. Two cytosines in CHG context and one in CHH context displayed minor methylation changes at 38°C, but these small fluctuations suggested no major regulatory remodeling **(Fig. 9c)**. This stability contrasted sharply with the dramatic shifts in the critical window, confirming that CritiCal-C specifically identifies environmentally responsive regulatory sites rather than constitutively methylated sequences. The experimental validation spanning transcriptional profiling, bisulfite sequencing of contrasting promoter regions, and correlation with expression changes establishes that CritiCal-C’s accurately captures the regulatory cytosines.

### The CritiCal-C resource portal and server

The findings made here have been converted into a first of its kind information and resource portal, CritiCal-C. CritiCal-C integrates data from in-house processed datasets, Plant Ensembl, UniProt, and KEGG database with advanced visualization method, allowing researchers to explore cytosine-specific effects on gene regulation. If a user wishes to explore critical cytosines in *A. thaliana* and *O. sativa* genes, the user can access the database, select the gene of interest, view complete GO details and its critical cytosines, and download the gene’s promoter sequence. For cross-species analysis, users can upload methylated 2kb promoter DNA sequences of their gene of interest in FASTA format. The back-end, leveraging a developed deep-learning model, processes the query sequences to estimate gene expression levels and identify critical cytosines. Using the same information results are displayed through interactive visualizations, with options for tabular downloads to support further analysis. Additionally, the web server provides Grad-CAM explainability scoring plots, highlighting the influence of each cytosine on the gene expression prediction. The overall system architecture and workflow are illustrated in **Supplementary File Materials and Methods, Fig. S8**, and detailed implementation information is provided in the **Supplementary File Materials and Methods**. The CritiCal-C database and web-server is freely accessible at: https://hichicob.ihbt.res.in/critical/

## Conclusion

Among all the four nucleotides of DNA, cytosine stands out for its stake in modulation of gene regulation, especially in plants. In plants, cytosine methylation is critical regulator of differential experession and guide the molecular system response with respect to environmental shifts. However, there is not much to identify such critical cytosines. The present study has attempted to fill in such gap to provide first ever such study, tool, and resource portal where criticality of cytosines has been measured using deep-learning. Critical cytosines exist as qualitative as well as quantitative ones. Experimental validation in *A. thaliana* under independent stress conditions confirms condition-specific methylation changes at identified regulatory critical cytosines accompanied by concordant transcriptional responses.

Homology based searches can’t identify critical cytosine across species, but GC% similarity explain it better. Using the developed tool here, CritiCal-C, it is now possible to identify critical cytosines in any species, besides successfully predicting the gene expression if bisulfite data is also given. This study will help to strategize molecular interventions like editing and targeted methylation/demethylation at critical cytosine for crop improvements and molecular studies, a way for minimal invasive and much easier molecular interventions.

## Acknowledgements

This study was conducted under the aegis of The Himalayan Center for High-throughput Computational Biology (HiCHiCoB), a BIC supported by DBT, Govt. of India. VK, AK, AS and SG are thankful to DBT, India for financial support as project associateship. KP acknowledges support from DBT-SRF. VK,SG and KP are thankful to Academy of Scientific and Innovative Research (AcSIR) for their Ph.D. enrollment. All authors are thankful to the Director, CSIR-IHBT, for his kind support for this study. This MS has CSIR-IHBT MSID 5832.

## Author’s contributions

VK performed a major part of this study. AK assisted in the development of the deep-learning model and developed the web server. AS developed the CritiCal-C database and implemented the front-end visualization. SG assisted with analysis evaluation. KP performed the wet-lab validation experiments under the guidance of GZ and prepared the corresponding validation section of the manuscript. RS conceptualized, designed, analyzed, and supervised the entire study. VK and RS wrote the manuscript.

## Supplementary Data statement

Supplementary Data are available at NAR online.

## Conflict of interest

The authors declare that they have no conflicts of interest.

## Funding

No funding received.

## Software and Data availability

The secondary data utilized in this study are publicly accessible, and their appropriate references and sources are listed in Supplementary File 2, Sheet 1-4. Additionally, the databse and server is available as a web server at https://hichicob.ihbt.res.in/critical/.

## Notes

### Competing Interest Statement

The authors have declared no competing interest.

### Summary of Updates

This revised version includes methodological clarification, improved experimental validation details, and enhanced figure presentation. The description of genome wide annotation of 2 kb promoter regions in Arabidopsis thaliana and rice has been refined for clarity and reproducibility. Experimental validation details have been expanded to better describe targeted bisulfite sequencing and qRT PCR under heat stress, including clearer explanation of locus selection and associated expression responses. Overstated language has been revised to maintain scientific neutrality and consistency throughout the manuscript. No major conclusions have changed. The revised manuscript improves methodological transparency, strengthens presentation of validation data, and provides clearer interpretation of results.

