## Supplementary File 1 for "Explainable Deep-Learning on condition specific expression profiles reveals critical cytosines in gene regulation"

**Table S1.** Comparison of DNA methylation detection methods.

| **Techniques** | **Aim** | **Strength** | **Weakness** | **References** |
|---|---|---|---|---|
| MSP | The candidate gene and/or the targeted CpGs are known | Cost-effective, Single-base resolution | Only few CpG sites can be detected through MSP, Low throughput | [1] |
| Pyrosequencing | The candidate gene and/or the targeted region are known | High quantitative and accurate resolution of DNA sequencing results through bioluminometric detection during nucleotide incorporation, Cost-effective, Single-base resolution | Degradation of DNA template due to bisulfite treatment, Validated primers are essential for the success of pyrosequencing are essential for the success of pyrosequencing | [2] |
| HM450K | De novo DNA methylation exploration | Cover most of CpG islands in human epigenome, Relatively cost-effective, Single-base resolution | Coverage is highly dependent on predesigned array | [3-5] |
| MRE-Seq | De novo DNA methylation exploration | Cost-effective, No harsh chemical treatment on DNA, thus avoid complicated experimental consequences | Coverage relies on restriction enzymatic activities and recognition sites, Only detect enrichment abundance, Can not identify individual CpG site, Can not detect absolute methylation level | [6] |
| MeDIP | De novo DNA methylation exploration | Cost-effective, No harsh chemical treatment on DNA, thus avoid complicated experimental consequences, Specific antibody against | Abundance resolution in ~100 bp, Can not identify individual CpG site, Can not detect absolute methylation level, The quality of the results relies on the quality of the antibody | [7] |

| | | | | |
|---|---|---|---|---|
| | | 5mC leads to sensitivity in regions with low CpG density | | |
| SMRT sequencing | De novo DNA methylation exploration | Ability to sequence native DNA through single molecule long-read sequencing, No harsh bisulfite treatment, Detect both nucleotide sequence and major types of DNA methylation patterns such as 5mC, 5hmC, 6mA and 4mC simultaneously, Particularly recommend to detect bacterial genomes | Because DNA cannot be amplified, large input DNA is required, Mostly applied in bacterial genome | [3, 8, 9] |
| OxBS-seq | De novo DNA methylation exploration for 5hmC | The most commonly used method to evaluate global 5hmC status | Multiple bisulfite treatments requires high amount of input DNA and high sequencing depths for confident determination of scarcely abundant modifications | [10] |
| PCR-based bisulfite sequencing | The candidate gene and/or the targeted region are known | Cost-effective, Allow detection of all CpG sites from the PCR products, Single-base resolution | Inconsistent results due to degradation of DNA template after bisulfite treatment, Require multiple replications (>10) to confirm the results | [11, 12] |
| WGBS | *De novo* DNA methylation exploration | The most comprehensive method to evaluate global DNA methylation state of almost every CpG site, | Expensive, Require sophisticated bioinformatics analysis, Require large amounts of input DNA due to degradation of DNA template after | [13, 14] |

| | | | | |
|---|---|---|---|---|
| | | Single-base resolution | harsh bisulfite treatment | |
| RRBS | De novo DNA methylation exploration | Cover the most representative CpG islands in the gene regulatory regions, Detect DNA methylation in different species, Single-base resolution, Relatively cost-effective | Only detect 1–3% of genome, May lose coverage at intergenic and distal regulatory regions | [15, 16] |
| ELISA-based assay | Broad prediction of global DNA methylation changes | Cost-effective, Commercially-available kit targeting 5mC | Only for rough estimation of global DNA methylation change, High variability and unreliable results | [17, 18] |
| Single-cell bisulfite sequencing | De novo DNA methylation exploration in single-cell level | Provide methylation information into individual cells, Particularly useful for specific cell types such as germ cells, embryonic stem cells, Have different sequencing options such as scWGBS and scRRBS | Same weaknesses in WGBS or RRBS assays | [19-21] |
| Nanopore sequencing | De novo DNA methylation exploration | Ability to sequence native DNA through single molecule long-read sequencing, No harsh bisulfite treatment, Applied in all species | Because DNA cannot be amplified, large input DNA is required, Lack of generalized algorithms makes it hard to understand stability of performance across species and sequencing batches | [3, 19, 22] |

**Table S2.** List of all terms involved in *A. thaliana* species

| Enrichment FDR | GO ID | GO Term |
|---|---|---|
| **Top 200** | | |
| **Biological Process** | | |
| 0 | GO:0043007 | Maintenance of rDNA |
| 0 | GO:0048658 | Anther wall tapetum development |
| 0 | GO:0009555 | Pollen development |
| 0 | GO:0048443 | Stamen development |
| **Cellular Component** | | |
| 0.01 | GO:1905369 | Endopeptidase complex |
| 0.01 | GO:0030176 | Integral component of endoplasmic reticulum membrane |
| 0.03 | GO:0009504 | Cell plate |
| 0.03 | GO:0005769 | Early endosome |
| 0.02 | GO:1905368 | Peptidase complex |
| 0.01 | GO:0005768 | Endosome |
| **Molecular Function** | | |
| 0 | GO:0042562 | Hormone binding |
| 0.01 | GO:0016307 | Phosphatidylinositol phosphate kinase activity |
| 0.01 | GO:0010329 | Auxin efflux transmembrane transporter activity |
| 0.01 | GO:0016740 | Transferase activity |
| **KEGG** | | |
| 0 | ath03015 | mRNA surveillance pathway |
| **Top 400** | | |
| **Biological Process** | | |
| 0 | GO:0010152 | Pollen maturation |
| 0 | GO:0048235 | Pollen sperm cell differentiation |

| | | |
|---|---|---|
| 0 | GO:0055046 | Microgametogenesis |
| 0 | GO:0048232 | Male gamete generation |
| 0 | GO:0022412 | Cellular proc. involved in reproduction in multicellular organism |
| 0 | GO:0007276 | Gamete generation |
| 0 | GO:0048555 | Cellular developmental proc. |
| **Cellular Component** | | |
| 0.04 | GO:0048555 | Generative cell nucleus |
| 0 | GO:0005887 | Integral component of plasma membrane |
| 0.01 | GO:0019005 | SCF ubiquitin ligase complex |
| 0.02 | GO:0031461 | cullin-RING ubiquitin ligase complex |
| 0.02 | GO:0140535 | Intracellular protein-containing complex |
| **Molecular Function** | | |
| 0 | GO:0030410 | Nicotianamine synthase activity |
| 0 | GO:0047498 | Calcium-dependent phospholipase A2 activity |
| 0 | GO:0010011 | Auxin binding |
| 0 | GO:0016165 | Linoleate 13S-lipoxygenase activity |
| 0 | GO:0052691 | UDP-arabinopyranose mutase activity |
| 0 | GO:0004623 | Phospholipase A2 activity |
| 0 | GO:0000285 | 1-phosphatidylinositol-3-phosphate 5-kinase activity |
| **KEGG** | | |
| 0 | ath00999 | Biosynthesis of various plant secondary metabolites |
| 0.04 | ath00592 | Alpha-Linolenic acid metabolism |
| 0.02 | ath04070 | Phosphatidylinositol signaling system |
| 0.02 | ath00562 | Inositol phosphate metabolism |
| 0.01 | ath00270 | Cysteine and methionine metabolism |

| | | |
|---|---|---|
| 0.03 | ath01250 | Biosynthesis of nucleotide sugars |
| 0.04 | ath00520 | Amino sugar and nucleotide sugar metabolism |
| 0.04 | ath01110 | Biosynthesis of secondary metabolites |
| 0.04 | ath01100 | Metabolic pathways |

**Table S3.** List of all terms involved in *O. sativa* species

| Enrichment FDR | GO ID | GO Term |
|---|---|---|
| **Top 200** | | |
| **Biological Process** | | |
| 0.01 | GO:0071071 | Phospholipid biosynthetic proc. |
| 0.01 | GO:1903725 | Phospholipid metabolic |
| 0.01 | GO:0019288 | Isopentenyl diphosphate biosynthetic |
| 0.01 | GO:0019682 | Glyceraldehyde-3-phosphate metabolic |
| 0.01 | GO:0019747 | Isoprenoid metabolic proc. |
| 0.01 | GO:0009240 | Isopentenyl diphosphate biosynthetic proc. |
| 0.01 | GO:0046490 | Isopentenyl diphosphate metabolic proc. |
| 0.01 | GO:0010675 | Cellular carbohydrate metabolic proc. |
| 0.01 | GO:0046890 | Lipid biosynthetic proc. |
| 0.01 | GO:0010027 | Thylakoid membrane organization |
| 0.01 | GO:0009668 | Plastid membrane organization |
| 0.01 | GO:0019216 | Lipid metabolic proc. |
| 0.02 | GO:0006081 | Cellular aldehyde metabolic proc. |
| 0.02 | GO:0062012 | Small molecule metabolic proc. |
| 0.02 | GO:0061077 | Chaperone-mediated protein folding |
| 0.03 | GO:0042180 | Cellular ketone metabolic proc. |
| 0.03 | GO:0006090 | Pyruvate metabolic proc. |
| 0.04 | GO:0051338 | Transferase activity |
| **Cellular Component** | | |
| 0.05 | GO:0009570 | Chloroplast stroma |
| 0.05 | GO:0009532 | Plastid stroma |

| | | |
|---|---|---|
| 0.02 | GO:0009507 | Chloroplast |
| 0.02 | GO:0009536 | Plastid |
| **Molecular Function** | | |
| 0.01 | GO:0004765 | Shikimate kinase activity |
| **KEGG** | | |
| 0.04 | osa00710 | Carbon fixation in photosynthetic organisms |
| 0.04 | osa00620 | Pyruvate metabolism |
| 0.04 | osa01200 | Carbon metabolism |
| **Top 400** | | |
| **Biological Process** | | |
| 0.02 | GO:0006357 | Cell wall macromolecule catabolic process |
| 0.02 | GO:0080022 | Aminoglycan catabolic process |
| 0.02 | GO:0010646 | Chitin metabolic process |
| 0.02 | GO:0048235 | Chitin catabolic process |
| 0.02 | GO:0009838 | Amino sugar catabolic process |
| 0.02 | GO:0009892 | Glucosamine-containing compound metabolic process |
| 0.02 | GO:0051094 | Glucosamine-containing compound catabolic process |
| 0.03 | GO:0010103 | Aminoglycan metabolic process |
| 0.03 | GO:0045892 | Amino sugar metabolic process |
| **Cellular Component** | | |
| 0 | GO:0033095 | aleurone grain |
| 0.04 | GO:0031410 | cytoplasmic vesicle |
| 0.04 | GO:0097708 | intracellular vesicle |
| 0.04 | GO:0031982 | vesicle |
| **Molecular Function** | | |
| 0.01 | GO:0045735 | nutrient reservoir activity |
| **KEGG** | | |
| 0.02 | osa00051 | Fructose and mannose metabolism |
| 0.02 | osa00710 | Carbon fixation in photosynthetic organisms |

| | | |
|---|---|---|
| 0.02 | osa01200 | Carbon metabolism |
