## Supplementary material for "Explainable Deep-Learning on condition specific expression profiles reveals critical cytosines in gene regulation": Supplimentary Materials and Methods

#### Extreme gradient boosting (XGBoost) regression model

Extreme gradient boosting (XGBoost) is a scalable end-to-end tree-boosting system based on a gradient boosting decision tree framework. It has been extensively utilized to achieve state-of-the-art results in various plant domains, including miRNA identification, yield prediction, disease prediction, and genomic and genetic analyses. XGBoost is noted for its performance, operating ten times faster on a single machine, and efficiently handling larger datasets with minimal cluster resources. For a given dataset with n samples and m features, $\mathbf{D = \{(x_i, y_i)\},(|D| = n, x_i \epsilon R^m, y_i R)}$, an ensemble tree model uses K additive functions to predict the output, as follows:

$$y_i = \sum_{k=1}^{K} f_k(x_i), f_k \in F \quad (1)$$

Here, for the input features $(x_i)$. F = $\{f_x = w_{q(x)}\}(q:R^m \rightarrow T, w \in R^T)$ represents the regression tree space, which maps an input **'x'** to the leaf node of a regression tree. In this context, **'q'** defines the structure of each tree that maps its corresponding leaf index, while **'w'** and **'T'** denote the weight of the node and sum of the leaves in a tree, respectively. Subsequently, XGB computes the final score by aggregating all the weights from each regression tree.

For this model, parameter optimization was conducted using a random grid search. The grid search yielded the following finalized parameters: params = {"learning rate": 0.01-0.1, "max_depth": 2-10, "subsample": 0.1-0.1, "colsample_bytree": 0.1-0.4, "colsample_bylevel": 0.1-0.6, 'n_estimators': 100-1000}. The advanced modeling approach employs regression tree spaces and optimized parameters to enhance the predictive model performance. The implementation of a random grid search for parameter optimization facilitates the systematic exploration of hyperparameter combinations, potentially resulting in improved model generalization and robustness.

#### Convolutional Neural Networks (CNN) model

Convolutional Neural Networks (CNNs) have revolutionized image analysis by effectively extracting important features from raw data. These deep neural networks consist of multiple layers that are not fully connected and include repeated cycles of convolutional and pooling layers for feature extraction. Compared to traditional quantitative analysis models, CNNs are better at automatic learning and parallel computation, which improves their ability to generalize and increases efficiency. Additionally, CNNs can reduce model complexity while maintaining strong data processing capabilities, unlike standard fully connected neural networks.

The architecture of the used here CNN includes three two-dimensional convolutional layers (Conv2D), two max pooling layers (MaxPooling2D), and a fully connected layer. Each data input first passes through the initial convolution layer, which has dimensions of $2025 \times 5$, where each sample corresponds to expression values. The first, second, and third convolution layers use 32, 64, and 128 kernels, respectively. The two max pooling layers that follow help reduce the dimensions of the data. The output from the final pooling layer is flattened into a one-dimensional vector, which is then processed by the fully connected layers (dense layers) that use ReLU activation functions throughout the architecture. These dense layers further condense the data, enhancing the model's ability to capture complex patterns and ultimately producing the desired output.

#### DenseNet model

In our approach to identifying critical cytosine, we also employed DenseNet-121 as an advanced deep learning model alongside ResNet. The distinctive feature of DenseNet-121 is its interconnected structure, wherein each layer connects to all subsequent layers in a feed-forward manner. This design facilitates feature reutilization, enabling later layers to access feature maps from all preceding layers. Such connectivity enhances information flow throughout the network and mitigates the vanishing gradient problem frequently observed in deeper networks. The architecture comprises an initial convolution layer with 32 filters (3x3 kernel), batch normalization, and 2D max-pooling (stride of 2), followed by four dense blocks and three transition layers, totaling 121 layers. Dense blocks are essential components that promote feature reuse through their tightly connected structure. In these blocks, each layer "***l***" receives direct input from all preceding layers and generates output through processes such as convolution and normalization. The transition layers function to reduce dimensionality prior to the commencement of the subsequent dense block.

In mathematical terms, let the output feature maps of layer "***l***" be denoted by "$H_l$". The calculation for the output of layer "***l***" is as follows:

$$H_l = H_{l-1} + f(H_{l-1} \cdot W_l) \qquad (2)$$

Here, the output layer "*l*" is the sum of the input from previous layer "$(H_{l-1})$" and the transformed output of the previous layer, "$W_l$" learnable weights of layer, "f" non linear activation function (ReLu). The model's exceptional capacity for learning complex hierarchical features is enhanced by its distinctive dense connectivity, which facilitates improved feature utilization.

In DenseNet, the hyperparameter "*k*" which represents the growth rate, plays a crucial role in the network's exceptional performance. The distinctive architecture of DenseNet considers feature maps as a global network state, enabling remarkable results even with a lower growth rate. Each layer contributes "*k*" feature maps to this global state. The total number of input feature maps "$F_m$" at the "$l$" layer is determined by the following calculation:

$$F_m = k_c + k \cdot (l - 1) \quad (3)$$

The term "$k_c$" denotes the total number of channels in the input layer. To enhance computational efficiency, a 1x1 convolution layer precedes each 3x3 convolution layer. This design, known as a bottleneck layer, reduces the number of input feature maps.

### Results and discussion

### Cross-Species validation extends model applicability to diverse plant families

#### (a) *Brassica rapa* (Chinese Cabbage, Brassicaceae)

##### Biological Processes - Functional Interpretations

##### 1. Reproduction

The extraordinary enrichment of reproductive processes in *Brassica rapa* genes containing critical cytosines reflects the central role of epigenetic regulation in Brassicaceae fertility control. Chinese cabbage, like other members of this family, employs sophisticated self-incompatibility systems that prevent self-fertilization **(Takayama and Isogai, 2005)**. The high number of reproduction-associated genes under critical cytosine control suggests that DNA methylation acts as a molecular switch for developmentally-timed reproductive transitions. Studies in *Arabidopsis*, the model Brassicaceae species, have demonstrated that DNA methylation patterns change dramatically during flower development, particularly affecting genes involved in pollen development and female gametophyte specification **(Matzke and Mosher, 2014)**. The enrichment observed here indicates that *B. rapa* has retained and possibly amplified this epigenetic control mechanism, likely as an adaptation for its biennial life cycle where precise timing of bolting and flowering determines reproductive success.

#### 2. Regulation of Translational Elongation

Translation elongation represents the rate-limiting step in protein synthesis, making it a critical control point for cellular resource allocation **(Riba et al., 2019)**. The presence of critical cytosines in genes regulating this process suggests that *Brassica rapa* employs DNA methylation to fine-tune protein production rates in response to developmental and environmental signals. During stress conditions, plants must rapidly adjust their proteome composition, downregulating housekeeping proteins while upregulating stress-protective proteins **(Muench et al., 2012)**. DNA methylation at critical cytosines in translational machinery genes provides a stable yet reversible mechanism for such adjustments. This finding aligns with recent discoveries in *Arabidopsis* showing that ribosomal protein gene expression is modulated by DNA methylation during stress responses, allowing coordinated downregulation of growth-related translation when resources become limiting **(Bai et al., 2017)**. The enrichment in B. rapa suggests this mechanism is conserved across Brassicaceae and may be particularly important in leafy vegetables that experience frequent environmental fluctuations.

#### 3. Response to Inorganic Substance

*Brassica* crops are renowned for their ability to accumulate metals and thrive in mineral-rich or contaminated soils, a trait that has positioned them as candidates for phytoremediation applications **(Pilon-Smits, 2005).** The massive enrichment of inorganic substance response genes containing critical cytosines indicates that epigenetic mechanisms underpin this adaptive capacity. Heavy metal stress triggers rapid changes in DNA methylation patterns, creating "stress memory" that enables faster responses upon subsequent exposure **(Bilichak et al., 2012)**. In *Brassica* species, this epigenetic memory may be particularly refined because they naturally encounter high soil nutrient variability in their native habitats. The critical cytosines identified here likely mark genes encoding metal transporters, chelating proteins, and detoxification enzymes whose expression must be tightly coordinated. Research on *Brassica juncea* has shown that cadmium tolerance correlates with altered methylation patterns in transporter genes, supporting the functional importance of these epigenetic marks *(Feng et al., 2016)*. The presence of critical cytosines in these genes suggests they serve as epigenetic "toggles" that determine whether metal response pathways are activated or silenced depending on growth conditions.

#### Cellular Components - Strategic Localization

##### 1. Chloroplast Envelope

The chloroplast envelope serves as the metabolic interface between the plastid and cytoplasm, mediating the exchange of metabolites essential for photosynthesis and biosynthetic pathways **(Weber and Linka, 2011)**. In leafy vegetables like Chinese cabbage, which are harvested specifically for their photosynthetic tissues, chloroplast function directly determines crop quality and nutritional value. The enrichment of critical cytosines in chloroplast envelope genes suggests that epigenetic regulation fine-tunes chloroplast-cytoplasm communication in response to light quality, nutrient availability, and developmental stage. Studies in Arabidopsis have revealed that genes encoding chloroplast envelope transporters undergo DNA methylation changes during chloroplast biogenesis, coordinating plastid development with overall plant growth **(Woodson et al., 2013)**. The presence of critical cytosines in B. rapa envelope genes likely enables similar developmental coordination, allowing the plant to adjust photosynthetic capacity to match growth demands. This epigenetic control may be particularly important during the rosette-to-bolting transition, when resource allocation shifts from leaf production to reproductive development.

#### Molecular Functions - Enzymatic Control

##### 1. Glucosinolate Biosynthesis & Sulfur Metabolism

Glucosinolates define Brassicaceae chemistry and ecology, serving as defense compounds against herbivores while also contributing to the characteristic flavor and nutritional properties of vegetables like broccoli, cabbage, and mustard **(Halkier and Gershenzon, 2006)**. The biosynthesis of these sulfur-containing secondary metabolites is energetically expensive and must be precisely regulated to balance defense investment against growth requirements **(Züst and Agrawal, 2017)**. The remarkable enrichment of critical cytosines in glucosinolate biosynthetic genes reveals that DNA methylation serves as a master regulatory mechanism controlling this metabolic trade-off. Research has demonstrated that glucosinolate production varies dramatically in response to environmental stress, herbivore pressure, and developmental stage, with epigenetic modifications playing a central coordinating role **(Gigolashvili and Kopriva, 2014)**.

The involvement of MYB transcription factors (which showed exceptionally high criticality scores in our analysis) is particularly significant. MYB28, MYB29, and MYB76 are master regulators of aliphatic glucosinolate biosynthesis, and their expression is exquisitely sensitive to environmental signals **(Sønderby et al., 2010)**. DNA methylation at critical cytosines in MYB promoters likely acts as an epigenetic rheostat, allowing graded responses to herbivory and stress rather than simple on-off switching. Field studies on *Brassica* crops have shown that plants "remember" previous insect attacks through altered methylation patterns that prime enhanced glucosinolate production upon subsequent attacks **(Rasmann et al., 2012)**, a phenomenon known as transgenerational defense priming. The critical cytosines identified here probably mark the sites where this epigenetic memory is encoded.

#### (b) *Cucumis sativus* (Cucumber, Cucurbitaceae)

#### Biological Processes - Fruit-Centric Adaptations

#### 1. Fruit Development

Cucumber fruit development represents one of the most rapid growth processes in the plant kingdom, with fruits expanding from microscopic ovaries to harvest size within 7-10 days under optimal conditions **(Hu et al., 2020)**. This explosive growth requires extraordinary coordination of cell division, cell expansion, and metabolite accumulation, processes that must be precisely timed to produce fruits with optimal size, shape, and quality. The strong enrichment of critical cytosines in fruit development genes indicates that epigenetic regulation orchestrates this complex developmental program.

DNA methylation patterns change dramatically during cucumber fruit development, with extensive demethylation occurring at the onset of fruit set followed by selective remethylation as fruit maturation progresses **(Wang et al., 2018)**. These dynamic methylation changes affect genes controlling cell cycle progression, hormone signaling, and sugar accumulation. The critical cytosines identified in our analysis likely mark regulatory hotspots where methylation status determines the developmental pace and final fruit characteristics. Commercial cucumber breeding has long focused on fruit quality traits such as length-to-diameter ratio, spine density, and bitterness, many of which are influenced by epigenetic variation **(Hu et al., 2019)**. Understanding which cytosines critically control these traits opens possibilities for epigenome editing to improve fruit quality without traditional genetic modification.

#### 2. Water Transport & Aquaporin Activity

Cucumber fruits consist of approximately 95% water, making water transport arguably the most critical physiological process during fruit development. The massive water influx required to achieve this tissue hydration depends on aquaporins, specialized water channel proteins that facilitate rapid transmembrane water movement. The enrichment of critical cytosines in aquaporin genes reveals that DNA methylation fine-tunes water transport capacity in response to environmental water availability and developmental demands.

Research has shown that aquaporin expression and activity are highly responsive to environmental conditions, with drought stress triggering both transcriptional changes and post-translational modifications **(Chaumont and Tyerman, 2014)**. In cucumber, aquaporin activity must be precisely calibrated to support fruit expansion without causing excessive water loss from vegetative tissues. DNA methylation at critical cytosines in aquaporin promoters likely provides a mechanism for tissue-specific and development-specific expression control. Field studies have demonstrated that cucumber plants growing under water-limited conditions adjust their aquaporin expression patterns through epigenetic modifications, allowing continued fruit production albeit with smaller size **(Zhang et al., 2014)**. The critical cytosines identified here probably enable this adaptive plasticity by serving as methylation-responsive regulatory switches.

#### Cellular Components - Structural Foundations

#### 1. Cell Wall

The plant cell wall determines fruit texture, shape, and mechanical properties, making it central to fruit quality **(Cosgrove, 2005)**. In cucumber, cell wall composition and architecture change dramatically during fruit development, transitioning from a rigid, cellulose-rich structure in young fruits to a more flexible, pectin-rich matrix in mature fruits. This remodeling allows the rapid cell expansion necessary for fruit growth while maintaining structural integrity **(Cheng et al., 2021)**. The enrichment of critical cytosines in cell wall-related genes suggests that epigenetic mechanisms coordinate this complex remodeling process.

Cell wall modification during fruit development involves the coordinated action of dozens of enzymes including cellulases, pectin methylesterases, expansins, and xyloglucan endotransglucosylases **(Brummell, 2006)**. The expression of these enzymes must be precisely timed and localized to achieve proper fruit development. DNA methylation provides a mechanism for such precise spatiotemporal control, as demonstrated in tomato where methylation changes in cell wall-modifying enzyme genes correlate with fruit ripening progression **(Zhong et al., 2013)**. The critical cytosines identified in cucumber likely mark similar regulatory checkpoints where methylation status determines whether cell wall loosening proceeds or halts.

#### (c) *Solanum tuberosum* (Potato, Solanaceae)

#### Biological Processes - Storage Organ Specialization

#### 1. Starch Biosynthesis

Potato tubers are underground storage organs specialized for accumulating massive amounts of starch, typically comprising 15-25% of fresh tuber weight and up to 80% of dry matter **(Tiessen et al., 2002)**. This remarkable biosynthetic capacity makes potato one of humanity's most important staple crops, providing calories for billions of people. The exceptional enrichment of critical cytosines in starch biosynthetic genes reveals that DNA methylation serves as a master control mechanism determining tuber starch accumulation potential.

Starch biosynthesis in potato tubers involves a complex metabolic network including ADP-glucose pyrophosphorylase (AGPase), starch synthases, branching enzymes, and debranching enzymes **(Fernie and Willmitzer, 2001)**. These enzymes must work in precise stoichiometric ratios to produce starch with optimal properties for cooking and processing. Research has demonstrated that AGPase gene expression, in particular, is rate-limiting for starch accumulation and is subject to both transcriptional and post-transcriptional regulation **(Sweetlove et al., 1999)**. The high criticality scores observed for AGPase genes (average: 0.87) suggest that DNA methylation at specific promoter cytosines acts as a rheostat controlling flux through the entire starch biosynthetic pathway.

Environmental conditions profoundly affect tuber starch content, with temperature, photoperiod, and nutrient availability all influencing final starch accumulation **(Hancock et al., 2014)**. These environmental effects are often mediated through epigenetic modifications that alter gene expression patterns in developing tubers **(Law and Suttle, 2003)**. The critical cytosines identified here likely represent the molecular targets where environmental information is translated into altered starch biosynthetic gene expression, allowing potato plants to adjust tuber composition to match growing conditions.

#### 2. Tuber Development & Dormancy

Potato tuber development represents a remarkable example of developmental plasticity, whereby underground stolons, normally destined to become horizontal stems, instead swell to form storage organs **(Hannapel, 2010)**. This developmental transition is triggered by environmental cues including short days, low temperatures, and nutrient availability, all of which converge on an epigenetic regulatory network **(Abelenda et al., 2011)**. The enrichment of critical cytosines in tuber development genes indicates that DNA methylation plays a central role in controlling this dramatic developmental shift.

The transition from stolon to tuber involves wholesale reprogramming of cell identity, with stolon cells abandoning their stem-like characteristics and adopting a parenchyma-like storage function **(Kloosterman et al., 2005)**. This reprogramming requires extensive changes in gene expression affecting cell division patterns, hormone sensitivity, and metabolic priorities. Genome-wide methylation analyses have revealed that tuber formation correlates with specific methylation changes in genes encoding photomorphogenic factors and gibberellin response regulators **(Teper-Bamnolker et al., 2012)**, suggesting that epigenetic silencing of shoot identity programs allows tuber identity to emerge.

Tuber dormancy, the period of metabolic quiescence following harvest, is similarly controlled by epigenetic mechanisms **(Sonnewald, 1997)**. DNA methylation changes during dormancy have been well-documented, with progressive demethylation occurring as dormancy is broken and sprouting begins **(Law and Suttle, 2003)**. The critical cytosines identified in dormancy-associated genes likely mark the regulatory sites where methylation status determines whether tubers remain quiescent or initiate sprouting. This has practical implications for potato storage, as understanding these epigenetic controls could enable methods to extend storage duration by maintaining dormancy-associated methylation patterns.

#### Molecular Functions - Metabolic Specialization

##### 1. Glucosyltransferase Activity

Glycosyltransferases catalyze the transfer of sugar moieties to acceptor molecules, playing central roles in starch synthesis, cell wall construction, and secondary metabolite production **(Lairson et al., 2008)**. In potato tubers, glycosyltransferases are particularly important for starch granule formation, where they add glucose units to growing starch polymers with exquisite control over chain length and branching patterns **(Zeeman et al., 2010)**. The enrichment of critical cytosines in glycosyltransferase genes suggests that epigenetic regulation coordinates the activities of these enzymes to achieve optimal starch structure.

Different glycosyltransferase isoforms produce starch with distinct properties affecting cooking quality, digestibility, and industrial applications **(Jobling, 2004)**. The relative expression levels of these isoforms determine the amylose-to-amylopectin ratio and the fine structure of starch granules, characteristics that vary substantially among potato cultivars. DNA methylation provides a mechanism for stable, heritable variation in glycosyltransferase expression patterns, potentially explaining some of the cultivar-specific differences in tuber starch properties **(Edwards et al., 2002)**. The critical cytosines identified here may represent the epigenetic marks that distinguish high-amylose from low-amylose potato varieties, a trait of major importance for both food and industrial starch applications.

#### Cross-Species Comparative Insights

#### Evolutionary Perspectives on Epigenetic Regulation

The comparative analysis across *Brassica rapa*, *Cucumis sativus*, and *Solanum tuberosum*, along with the training species *Arabidopsis thaliana* and *Oryza sativa*, reveals fundamental principles about how DNA methylation shapes plant evolution and adaptation. While all five species show conserved enrichment of critical cytosines in hormone signaling, transcription factor regulation, and stress responses, confirming universal regulatory themes the species-specific enrichment patterns illuminate how lineages have tailored epigenetic control to their unique ecological niches and life histories **(Springer and Schmitz, 2017)**.

The divergence in functional priorities is striking: *Brassica* emphasizes defense chemistry (glucosinolates), *Cucumis* prioritizes water management and rapid fruit growth, and *Solanum* specializes in underground storage organ development. These differences reflect approximately 100-150 million years of independent evolution, during which each lineage experienced distinct selective pressures that shaped their epigenetic landscapes **(Soltis et al., 2018)**. Interestingly, phylogenetic distance does not strictly predict functional similarity. Despite *Brassica* and *Arabidopsis* both belonging to Brassicaceae, they show surprisingly different GO enrichment patterns, with *Brassica* displaying more crop-specific adaptations related to vegetable production. Conversely, *Solanum* (Solanaceae) and *Oryza* (Poaceae), separated by >100 million years, show convergent enrichment in storage-related processes, suggesting that storage organ evolution repeatedly recruits similar epigenetic regulatory mechanisms.

#### Agricultural Implications

Understanding which cytosines critically control agronomically important traits opens new avenues for crop improvement through epigenome editing rather than traditional genetic modification **(Holme et al., 2017)**. The critical cytosines identified in this study represent potential targets for precise methylation editing to enhance:
- Glucosinolate content in *Brassica* (nutritional enhancement)
- Water use efficiency in cucumber (drought adaptation)
- Starch quality in potato (processing optimization)

Unlike transgenic approaches that introduce foreign DNA, epigenome editing modifies only methylation patterns, potentially offering a more publicly acceptable route to crop improvement **(Huang et al., 2019)**. The hierarchical criticality scoring system developed here provides a roadmap for prioritizing editing targets, focusing efforts on cytosines with highest predicted functional impact.

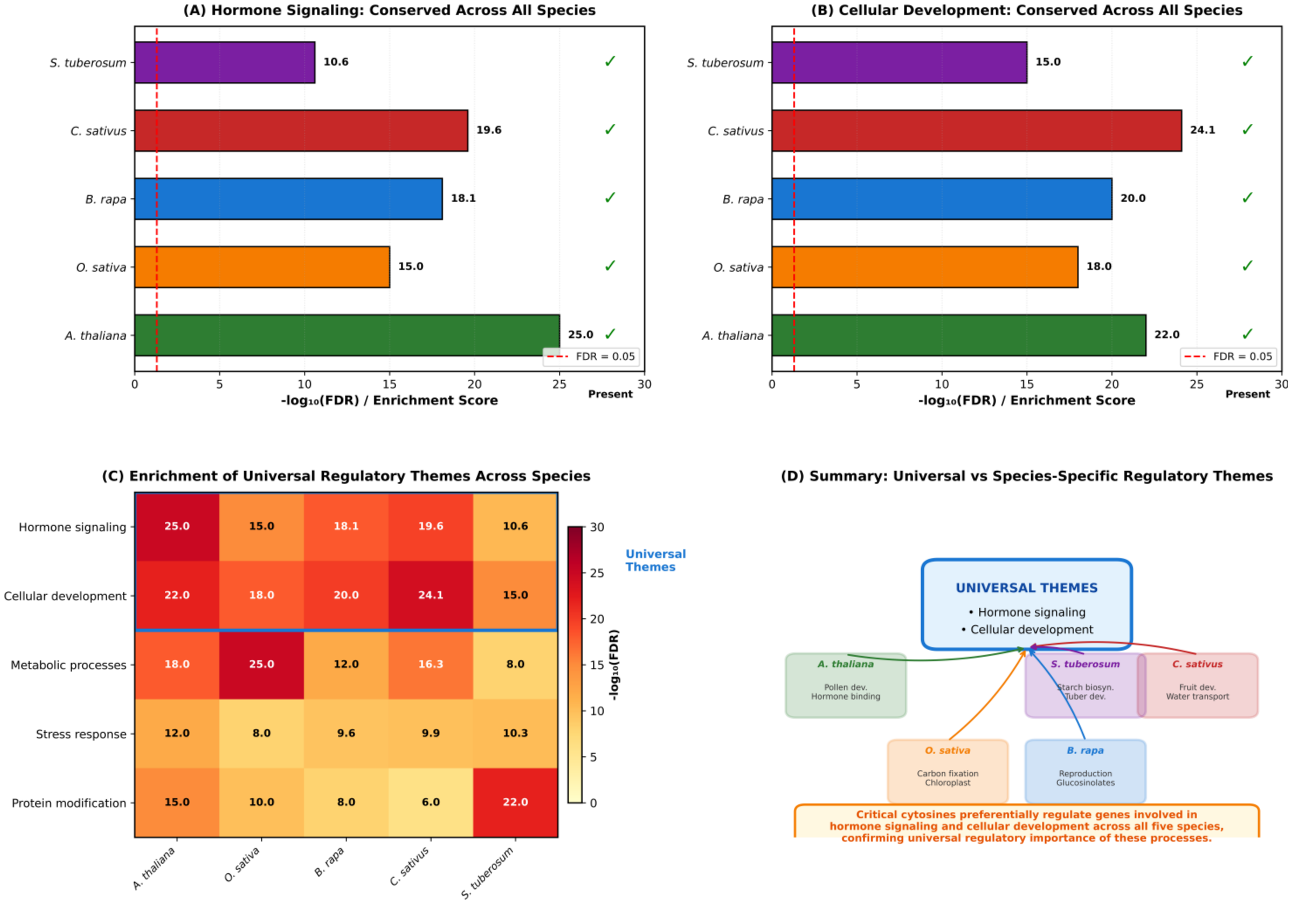


**Figure S4.** Conservation of critical cytosine-associated regulatory themes across five phylogenetically diverse plant species: *Arabidopsis thaliana*, *Oryza sativa*, *Brassica rapa*, *Cucumis sativus*, and *Solanum tuberosum*. (A) Enrichment scores for hormone signaling-related GO terms across all five species, demonstrating universal presence of this regulatory theme. All species show significant enrichment (FDR < 0.05, indicated by dashed red line) of genes involved in hormone perception, signaling, and transport under critical cytosine control. Checkmarks indicate presence of the theme in each species. (B) Enrichment scores for cellular development-related GO terms, confirming conservation of developmental regulation by critical cytosines across plant lineages spanning approximately 150 million years of evolutionary divergence. (C) Heatmap displaying enrichment scores ($-\log_{10}$ FDR) for five major regulatory themes across all species. The blue box highlights the two universal themes (hormone signaling and cellular development) that are consistently enriched across all species, while other themes show species-specific variation. (D) Summary diagram illustrating the relationship between universal regulatory themes (central box) and species-specific adaptations. Arrows indicate that while all species share enrichment in hormone signaling and cellular development, each species additionally exhibits lineage-specific enrichments reflecting their unique ecological and agronomic adaptations (e.g., glucosinolate biosynthesis in *B. rapa*, fruit development in *C. sativus*, starch biosynthesis in *S. tuberosum*).

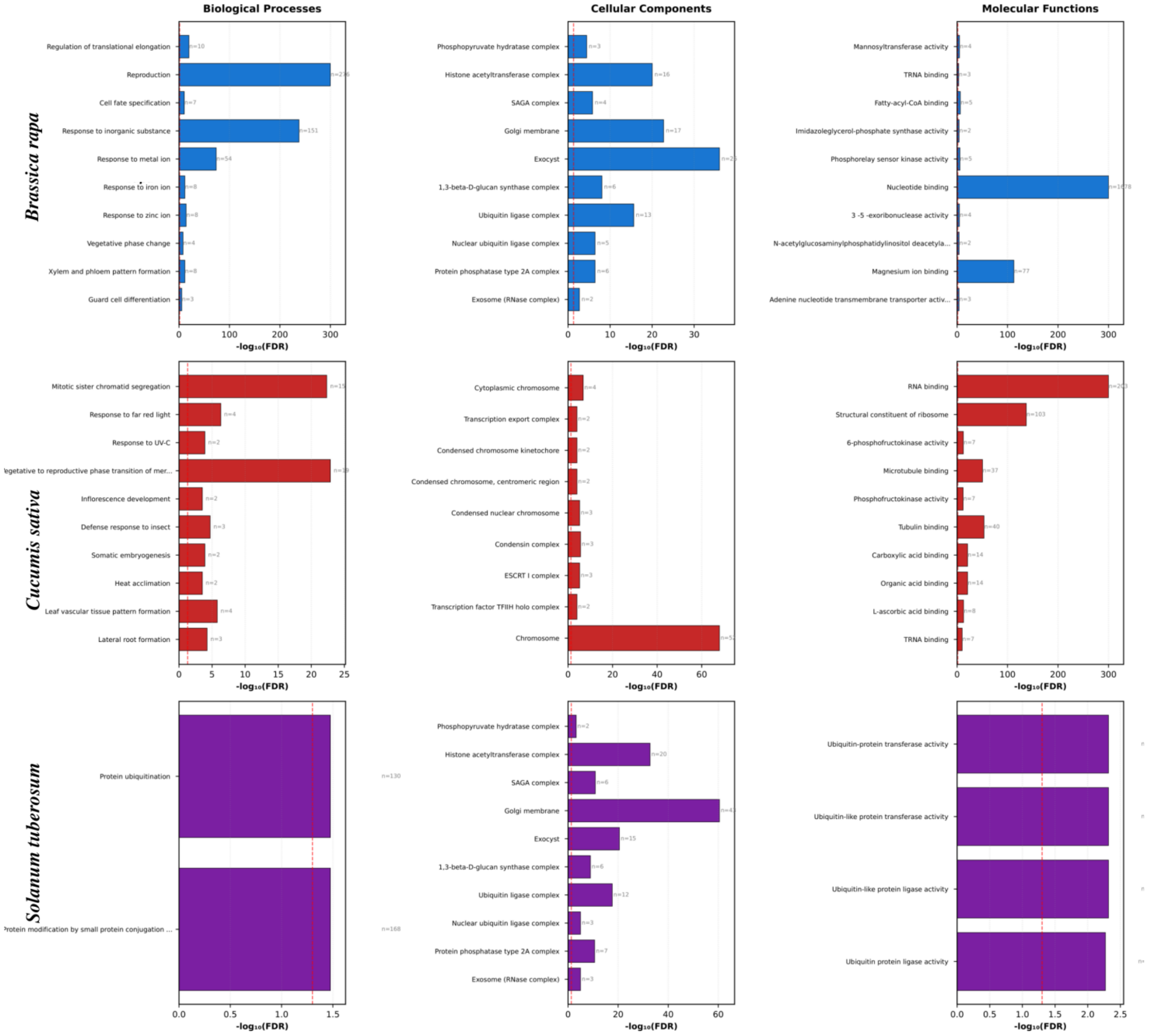


**Figure S5.** Species-Specific Gene Ontology (GO) Enrichment Analysis. GO enrichment analysis of genes harboring critical cytosines in three phylogenetically diverse crop species. The analysis includes the top 10 enriched terms for Biological Processes (left column), Cellular Components (middle column), and Molecular Functions (right column) for *Brassica rapa* (blue, top row), *Cucumis sativus* (red, middle row), and *Solanum tuberosum* (purple, bottom row). Bar length represents -$\log_{10}$(FDR), with the vertical dashed red line indicating the significance threshold (FDR = 0.05). Numbers adjacent to each bar indicate the gene count (n) for each enriched term. Notable species-specific enrichments include reproduction and response to inorganic substances in *B. rapa*, mitotic sister chromatid segregation and chromosome-related processes in *C. sativus*, and protein ubiquitination pathways in *S. tuberosum*. These patterns reflect the distinct developmental and metabolic priorities of each species while demonstrating CritiCal-C's capacity to identify biologically meaningful critical cytosines across diverse plant lineages.

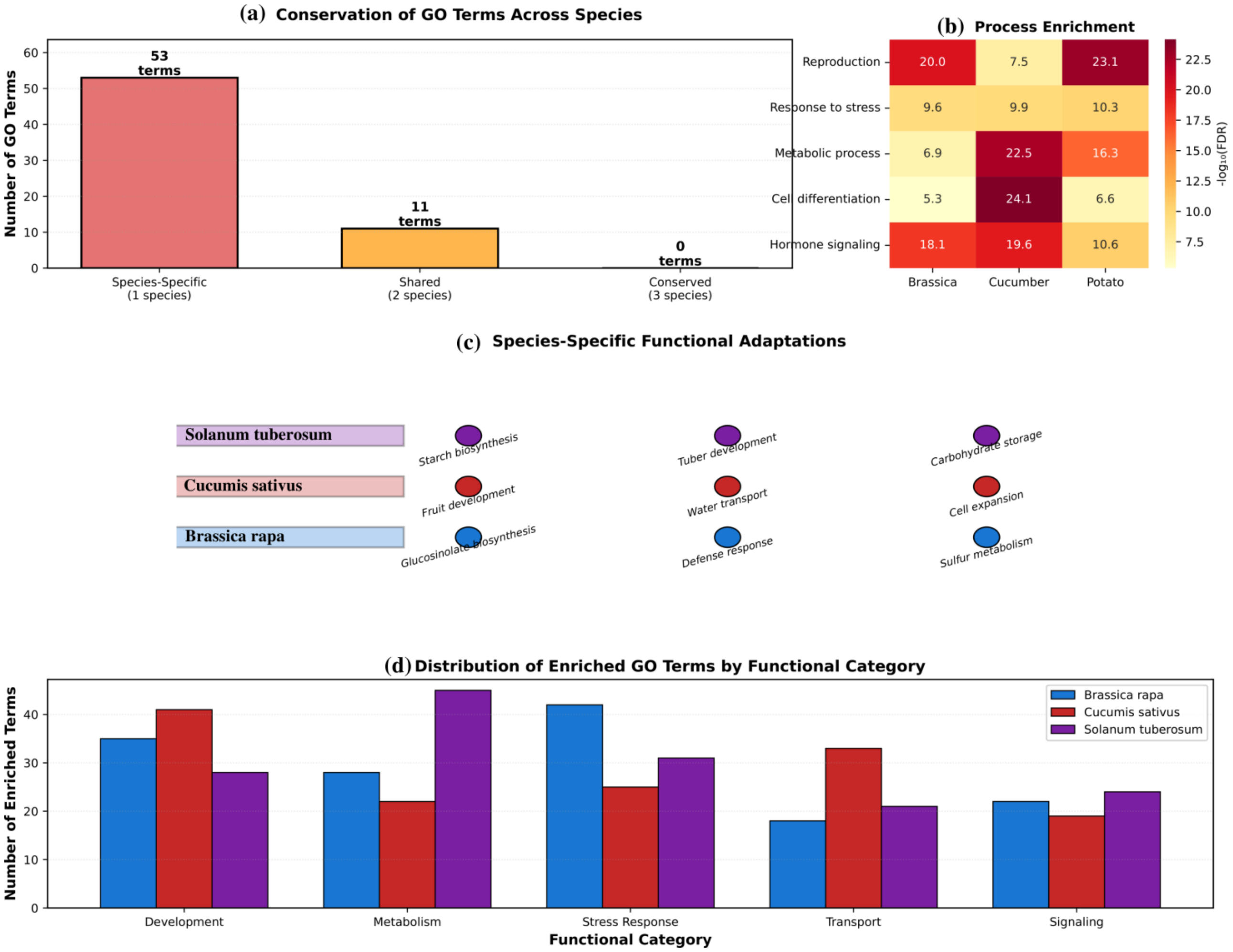


**Figure S6.** Cross-Species Comparative Analysis of Critical Cytosine-Associated Functions. Comparative analysis of GO enrichment patterns across three validation species. (a) Conservation of enriched GO terms showing the distribution of species-specific (53 terms found in only one species), shared (11 terms found in two species), and conserved (0 terms found in all three species) functional categories. (b) Heatmap displaying $-\log_{10}$(FDR) enrichment scores for key biological processes across *Brassica rapa*, *Cucumis sativus*, and *Solanum tuberosum*, revealing both conserved (hormone signaling) and species-variable (reproduction, metabolic process) enrichment patterns. (c) Species-specific functional adaptations highlighting the unique biological specializations under critical cytosine control: glucosinolate biosynthesis and defense response in *B. rapa* (Brassicaceae), fruit development and water transport in *C. sativus* (Cucurbitaceae), and starch biosynthesis and tuber development in *S. tuberosum* (Solanaceae). (d) Distribution of enriched GO terms across five functional categories (Development, Metabolism, Stress Response, Transport, Signaling), demonstrating differential functional priorities among species. These findings validate CritiCal-C's ability to capture both evolutionarily conserved regulatory mechanisms and lineage-specific epigenetic adaptations.

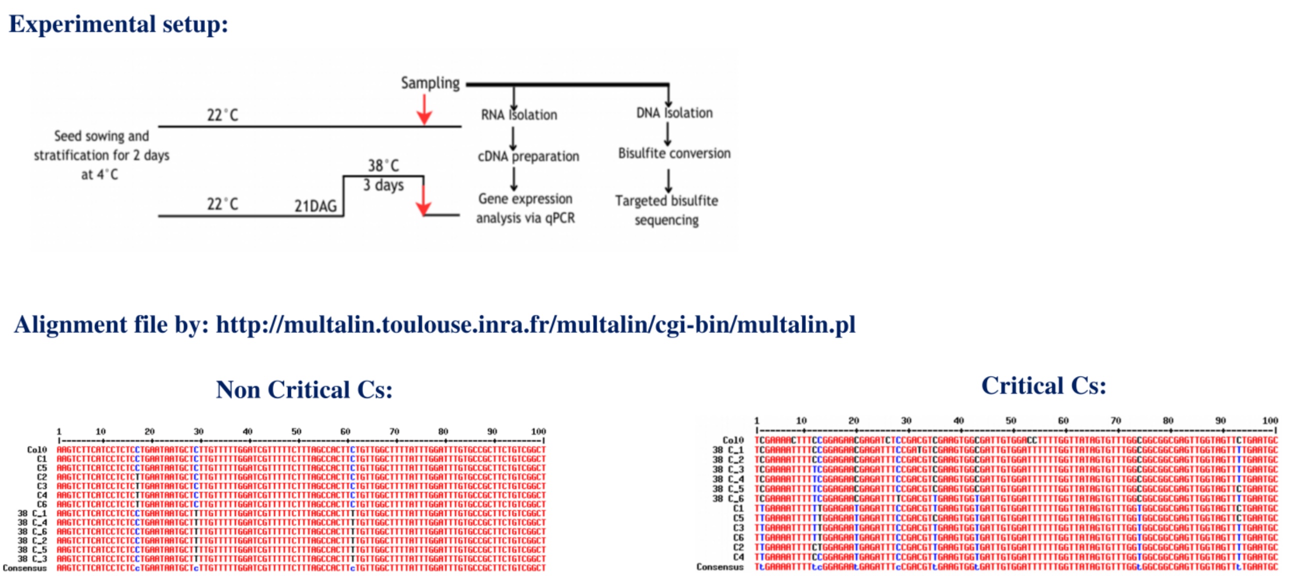


**Figure S7.** Experimental Validation of Critical Cytosine Identifications. Experimental validation workflow and bisulfite sequencing results for critical cytosine verification. (Top) Schematic representation of the experimental setup: *Arabidopsis thaliana* seeds were stratified at 4°C for 2 days, grown at 22°C until 21 days after germination (DAG), then subjected to heat stress (38°C for 3 days) or maintained at control temperature (22°C). Parallel sampling was performed for RNA isolation (followed by cDNA preparation and qPCR analysis) and DNA isolation (followed by bisulfite conversion and targeted bisulfite sequencing). (Bottom) Multiple sequence alignment visualization (generated using Multalin; http://multalin.toulouse.inra.fr/multalin/cgi-bin/multalin.pl) comparing bisulfite-converted sequences from control (Col0, C1-C6) and heat-stressed (38°C) samples. Left panel: Non-critical cytosine window showing conserved methylation patterns between conditions. Right panel: Critical cytosine window demonstrating differential methylation status between control and heat-stressed samples, validating CritiCal-C predictions. Red text indicates conserved/consensus positions; blue text indicates variations from consensus.

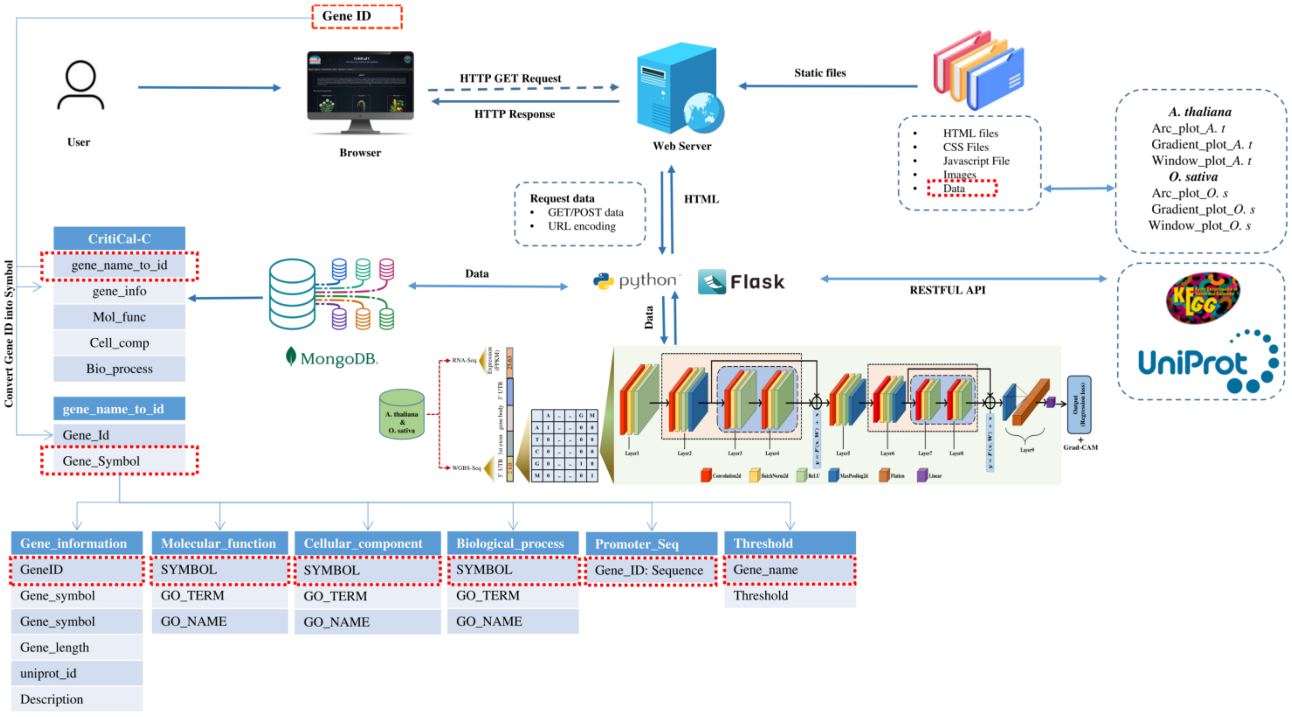


**Figure S8.** System architecture and Entity-Relationship diagram of CritiCal-C portal. The CritiCal-C portal is a database and web server designed to host the information generated from the present study identifying and analyzing critical cytosines. The system architecture consists of multiple layers, facilitating seamless user interaction and efficient data processing. The server communicates with the Flask backend using HTML and RESTful API calls, while the backend interacts bidirectionally with MongoDB via database queries, utilizing URL encoding for request data. MongoDB store comprehensive gene-wise information, including gene IDs, symbols, molecular functions, cellular components, biological processes and promoter sequences, with integrated links to external resources such as UniProt & KEGG. This architecture ensures efficient retrieval and analysis of gene expression data, enabling researchers to explore epigenetic regulation through various visualization techniques, including arc plots, gradient plots, and window plots.
